# Antagonist actions of CMK-1/CaMKI and TAX-6/Calcineurin along the *C. elegans* thermal avoidance circuit orchestrate adaptation of nociceptive response to repeated stimuli

**DOI:** 10.1101/2024.09.18.613419

**Authors:** Martina Rudgalvyte, Zehan Hu, Dieter Kressler, Joern Dengjel, Dominique A. Glauser

**Affiliations:** Department of Biology, University of Fribourg, 1700 Fribourg, Switzerland; Metabolomics and Proteomics Platform (MAPP), Department of Biology, University of Fribourg, 1700 Fribourg, Switzerland

## Abstract

Response decrease following repeated exposure to innocuous or noxious stimuli is a conserved adaptation phenomenon often referred to as habituation. Impaired nociceptive habituation is associated with several pain conditions in human, but the underpinning molecular mechanisms are only partially understood. In the nematode *Caenorhabditis elegans*, thermo-nociceptive adaptation to repeated stimuli was previously shown to be regulated by the Ca^2+^/Calmodulin-dependent protein kinase 1 (named CMK-1), but its downstream effectors were unknown. Here, using *in vitro* kinase assays coupled with mass-spectrometry-based phosphoproteomics, we empirically identified hundreds of CMK-1 phospho-substrates. Among them, we found that CMK-1 can phosphorylate the calcineurin A (CnA) protein TAX-6 *in vitro* in a highly conserved regulatory domain, which led us to hypothesize that TAX-6/CnA might be a downstream mediator of CMK-1 signaling in the control of thermo-nociceptive adaptation. Combined genetic and pharmacological manipulations revealed a network of antagonistic actions between CMK-1 and calcineurin pathways in the regulation of the responsiveness of naïve worms and the response adaptation to repeated noxious heat stimuli. However, the results of cell-specific rescue and gain-of-function experiments suggested that CMK-1 acts in AFD and ASER thermosensory neurons and that TAX-6/CnA acts in FLP thermosensory neuron and a set of downstream interneurons to regulate noxious heat avoidance behaviors. Because CMK-1 and TAX-6/CnA act in non-overlapping cell types, the phosphorylation event identified *in vitro* might not be relevant for this phenotype and the complex interaction between the two pathways might rather originate from their action in separate parts of the nervous system. As a whole, our study has identified (i) CMK-1 substrate candidates, which will fuel further research on the intracellular actuation of CMK-1-dependent signaling on various processes, and (ii) a complex set of antagonistic interactions between CMK-1 and calcineurin signaling operating at distributed loci within a sensory-behavior circuit, acting to adjust baseline thermo-nociception and regulate thermo-nociceptive adaptation.

## INTRODUCTION

Animals from invertebrates to mammals can detect and avoid harmful stimuli, which is essential for their survival and well-being. Noxious stimuli are detected and encoded by specialized primary sensory neurons in a process called nociception (Dubin and Patapoutian 2010). Nociceptive signals are relayed in the nervous system via synaptic connections to be further processed and trigger different responses, such as behavioral changes and the perception of pain (Treede 1995). Pain sensitivity is not fixed and can be adjusted in various physiological and pathological contexts through nociceptive plasticity processes such as hyperalgesia or nociceptive desensitization/habituation (Kidd and Urban 2001, Sandkühler 2009, Rodriguez-Raecke, Doganci et al. 2010, Breimhorst, Hondrich et al. 2012, May, Rodriguez-Raecke et al. 2012, O’Neill, Brock et al. 2012, Basith, Cui et al. 2016). Prolonged or repeated exposure to noxious stimuli can lead to either sensitization or desensitization/habituation depending on the stimuli intensity, frequency and various physiological factors (Treede 1995). Because impaired nociceptive habituation is linked to various human chronic pain conditions (de Tommaso, Lo Sito et al. 2005, Smith, Tooley et al. 2008, de Tommaso, Federici et al. 2011), obtaining a deeper understanding of the cellular and molecular processes into play appears highly relevant to aid in the development of new therapeutical strategies for pain management. Several molecular mechanisms have been shown to modulate nociceptive pathways from the periphery to the brain (Kuner and Flor 2017, Pace, Passavanti et al. 2018, Amna, Salman et al. 2019), which involve the regulated activity of proteins such as membrane receptors or ion channels (involved in sensory-transduction, cell excitability, or signal conduction). Many kinases and phosphatases, which actuate protein regulation via the control of their phosphorylation status and that were previously shown to broadly regulate neural plasticity in the nervous system, are also involved in nociceptive plasticity (Willis Jr. 2001, Fu, Han et al. 2008, Wang and Zhang 2012, Isensee, Diskar et al. 2014, Pace, Passavanti et al. 2018). These regulatory pathways include intracellular signaling by calcium/calmodulin-dependent protein kinases (CaMKs) (Shum, Ko et al. 2005, Liang, Liu et al. 2012, Schild, Zbinden et al. 2014, Zhou, Liu et al. 2017), which might couple cellular activity levels with pleiotropic intracellular effects via post-translational modifications over a large repertoire of potential target proteins. Identifying the phosphorylation substrates of these kinases that act in the nociceptive pathway to mediate plasticity effects could provide relevant insight for future pain management translational research.

The nematode *Caenorhabditis elegans* has emerged as a powerful model to study the molecular and cellular bases of nociception and its plasticity. *C. elegans* produces innate avoidance behaviors in response to a variety of noxious stimuli, including irritant chemicals, harsh touch and noxious heat (Kaplan and Horvitz 1993, Wittenburg and Baumeister 1999, Hilliard, Apicella et al. 2005, Li, Kang et al. 2011, Liu, Schulze et al. 2012). Noxious heat stimuli targeting the animal head or the entire animal trigger a stereotyped reversal behavior (Wittenburg and Baumeister 1999, Liu, Schulze et al. 2012, Byrne Rodgers and Ryu 2020, Lia and Glauser 2020), which involves several thermo-sensory neurons, including AFD, AWC, and FLP, proposed to work as thermo-nociceptors (Chatzigeorgiou and Schafer 2011, Liu, Schulze et al. 2012, Kotera, Tran et al. 2016). The molecular components underpinning the function of the nociceptive system are well-conserved. E.g., transient receptor potential (TRP) channels mediate worm thermo-nociceptive responses, like in fly and mammals (Chatzigeorgiou, Yoo et al. 2010, Glauser, Chen et al. 2011, Liu, Schulze et al. 2012, Nkambeu, Salem et al. 2020). Furthermore, persistent or repeated noxious heat stimuli cause a progressive reduction in the rate of heat-evoked reversals (Lia and Glauser 2020, Jordan and Glauser 2023), which we will refer to here as thermo-nociceptive adaptation. A genetic screen for human pain-associated gene orthologs revealed thermo-nociceptive adaptation alterations for many corresponding worm mutants, substantiating the interest of the model (Jordan and Glauser 2023). Among conserved molecular players, the worm CaMKI (named CMK-1) was shown to mediate thermo-nociceptive adaptation (Schild, Zbinden et al. 2014, Lia and Glauser 2020, Jordan and Glauser 2023). CMK-1 is broadly expressed in the worm nervous system and, beyond thermo-nociceptive adaptation, controls multiple experience-dependent plasticity processes, such as those related to developmental trajectories (Neal, Takeishi et al. 2015), the encoding of preferred growth temperature (Yu, Bell et al. 2014), salt aversive learning (Lim, Fehlauer et al. 2018) and habituation to repeated touch stimuli (Ardiel, McDiarmid et al. 2018). CMK-1 was shown to regulate the expression of the *guanylyl cyclase-8* (*gcy-8*) gene (Satterlee, Ryu et al. 2004), the AMPA glutamate receptor-1 (*glr-1)* gene (Moss, Park et al. 2016), the ortholog of the human DACH1/2 Dachsund transcription factor gene *(dac-1)* and the PY domain transmembrane protein-1 (*pyt-1*) gene (Harris, Bates et al. 2023). The impact of CMK-1 on gene transcription is partially mediated by the CREB homolog-1 (CRH-1) transcription factor (Harris, Bates et al. 2023), which could be phosphorylated by CMK-1 *in vitro* (Kimura, Corcoran et al. 2002) and in an heterologous expression system (Eto, Takahashi et al. 1999). A previous study used an *in silico* approach to systematically predict potential CMK-1 phosphorylation substrates and revealed repeated-touch habituation phenotypes in mutants for several candidate targets (Ardiel, McDiarmid et al. 2018). Apart from these few studies, we still know very little on the CMK-1 downstream effectors that mediate the numerous biological actions of CMK-1, including in mediating thermo-nociceptive adaptation.

Calcineurin is a well-conserved eukaryotic Ca^2+^/Calmodulin activated Serine/Threonine phosphatase (Klee, Crouch et al. 1979, Rusnak and Mertz 2000), involved in Ca^2+^ signaling pathway and functioning in several tissues, such as muscles, T cells and neurons, where it regulates synaptic plasticity and memory (Groth, Dunbar et al. 2003, Mukherjee and Soto 2011). Calcineurin regulates the activity of numerous proteins, such as transcription factors (Molkentin 2004), receptors, mitochondria proteins and microtubules depending on Ca^2+^ signaling state (Peuker, Strigli et al. 2022). Overexpression of calcineurin can cause cardiac hypertrophy (Gillis, Kavanagh et al. 2004, Yuan, Shen et al. 2024), higher infarct volumes (Dai, Hu et al. 2024), whereas lack of function can cause defects in kidney development (Gooch, Toro et al. 2004), and increase in autophagy (Ke, Huang et al. 2023). Chronic inhibition of calcineurin with immunosuppressant drugs, like cyclosporin A, was shown to cause irreversible neuropathic pain in some patients, a phenomenon referred to as calcineurin inhibitor-induced pain syndrome (Smith 2009). Several studies in mice have highlighted that calcineurin signaling modulate thermal nociception (Sato, Onaka et al. 2007). Several pathways working downstream of calcineurin signaling have been proposed to mediate the regulation of nociception, including NFAT transcription factors, TWIK-related spinal cord potassium channels (TRESK), TRP channels, and α2δ-1 Ca^2+^ channels (Smith 2009, Huang, Chen et al. 2020, Huang, Chen et al. 2022). Nevertheless, we still have a cursory understanding of how calcineurin signaling integrates with other intracellular signaling pathways at different locus in the nociceptive circuit to orchestrate nociceptive plasticity.

Calcineurin is a heterodimeric protein consisting of one calcineurin A (CnA) catalytic subunit, and one calcineurin B (CnB) regulatory subunit. CnA contains catalytic and autoinhibitory domains, as well as CnB and Ca^2+^/Calmodulin binding domains. In the absence of Ca^2+^ signaling, the autoinhibitory domain suppresses catalytic phosphatase activity. Increase in cytosolic Ca^2+^ promotes Ca^2+^ binding to CnB and to CaM. Ca^2+^/CnB and Ca^2+^/CaM in turn bind to calcineurin A subunit to release the autoinhibition, allow substrates to access the active site and thereby the activation of CnA (Bandyopadhyay, Lee et al. 2002, Kuhara, Inada et al. 2002, Wang, Du et al. 2008). *Caenorhabditis elegans* genome encodes one CnA homolog *tax-6 (*aka *cna-1)*, and one CnB homolog *cnb-1* displaying conserved structural and biochemical features with vertebrate calcineurin proteins (Bandyopadhyay, Lee et al. 2002, Bandyopadhyay, Lee et al. 2004). Lack of *tax-6* function causes thermal hypersensitivity and increased osmosensation, as well as increased adaptation of olfactory system (Kuhara, Inada et al. 2002). Loss-of-function mutants display a thermophilic behavior, and this anomaly could be rescued by wild type *tax-6* expression in AFD neurons. *tax-6* gain-of-function mutants show defective enteric muscle contraction (Lee, Song et al. 2005), as well as higher brood size and hypersensitivity to serotonin (Lee, Jee et al. 2004). The loss of *cnb-1* causes lethargic movements, delayed egg laying and reduced growth (Bandyopadhyay, Lee et al. 2002).

Here, we combined *in vitro* kinase assays with shotgun phosphoproteomics to empirically identify CMK-1 phospho-substrates in worm peptide and protein libraries. CMK-1 was found to phosphorylate TAX-6/CnA in a highly conserved regulatory region, suggesting cross-regulation between CaMK and calcineurin signaling in worms. We next combined genetic and pharmacological manipulations with a quantification of noxious heat reversals to assess the role of CMK-1 and TAX-6 signaling and their interactions in controlling naïve animal responsiveness and the adaptation to repeated stimulation. These follow-up analyses confirmed a complex set of antagonistic signaling actions produced by the two pathways. However, the two intracellular signaling pathways appear to primarily function in separate neuron types to control thermo-nociceptive adaptation indicating that the phosphorylation of TAX-6/CnA by CMK-1 might not be relevant *in vivo* for the control of thermo-nociception. Collectively, our results reveal multiple direct and indirect interactions between CaMK and calcineurin signaling and pave the role for deeper studies on the molecular control of nociceptive adaptation in a simple genetic model.

## RESULTS

### CMK-1 phosphorylates multiple substrates *in vitro* including TAX-6/CnA

To identify phosphorylation substrates of CMK-1, we performed *in vitro* kinase assays on two types of substrate libraries followed by mass spectrometry-based phosphoproteomics (Fig. 1A). To that end, we purified recombinant CMK-1(1-295)T179D mutant protein produced in *E. coli.* This mutant is a constitutively active form of CMK-1 that lacks the C-terminal auto-inhibitory domain and harbors a T179D phosphomimic mutation, thus bypassing the need for Ca^2+^/CaM binding and for phosphorylation by CKK-1. In two separate experiments, we assessed the phosphorylation produced by CMK-1(1-295)T179D, first, on a library of worm peptides produced by trypsin digestion of whole protein extracts and, second, on a whole protein library produced in non-denaturing conditions. As expected given the strong structural and functional conservation within the CaM kinase family, the consensus substrate motifs obtained with each substrate libraries matched the ΦXRXX(S/T)XXXΦ consensus previously characterized for mammalian CaMK (where Φ represents hydrophobic residues, Fig. 1B) (Lee, Kwon et al. 1994, White, Kwon et al. 1998). This result supports the validity of our *in vitro* kinase assays to identify direct CMK-1 targets and indicates that our experimental design efficiently mitigated the impact of any potential kinase activity coming from *E. coli* protein contamination or from worm kinases in the case of whole protein extracts.

**Figure 1.**
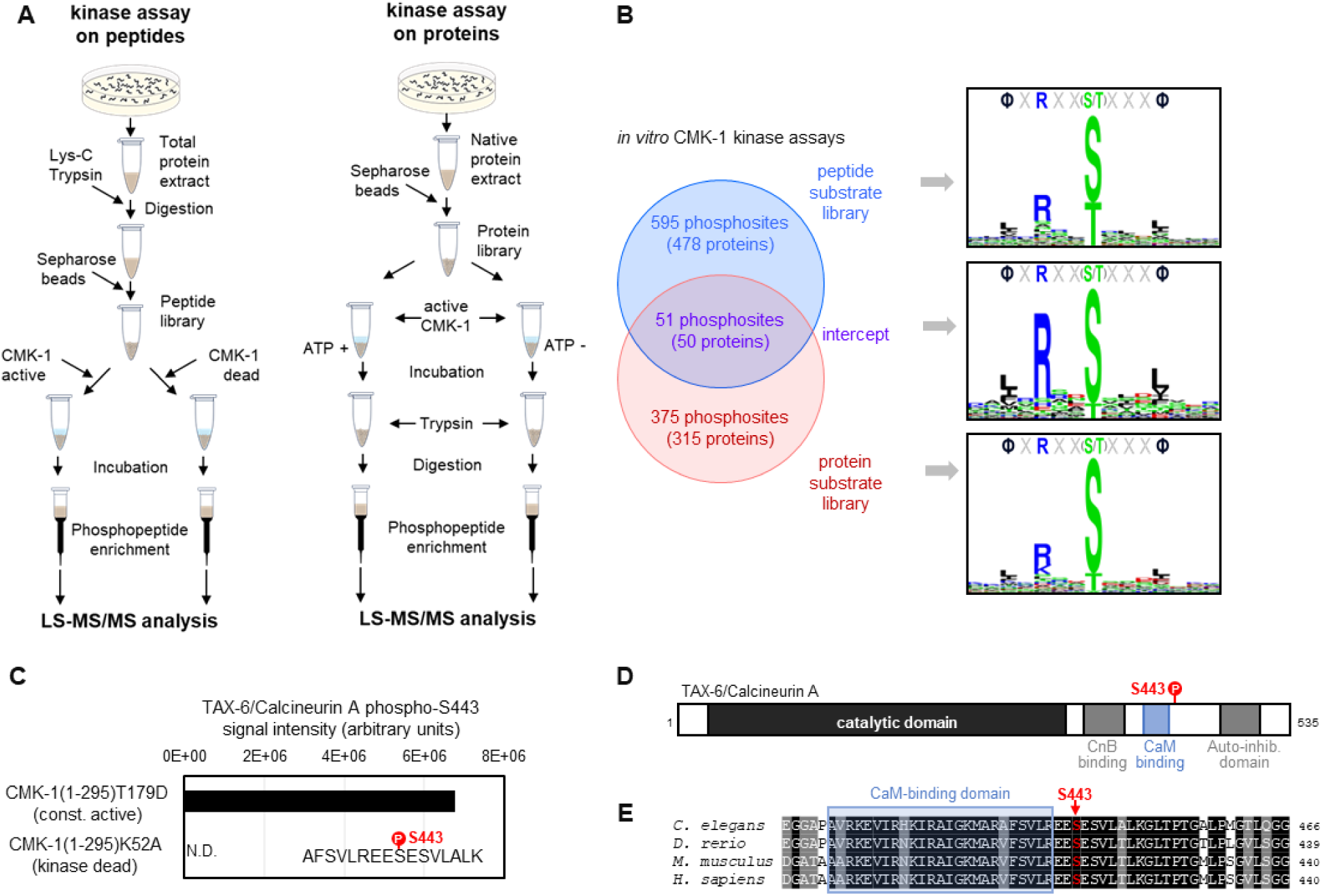
Identification of CMK-1 phosphorylation substrates. **A**. Schematic of the two approaches used for the large-scale identification of the CMK-1 phosphorlyation substrates *in vitro*. *In vitro* kinase assays using a *C. elegans* peptide library (left) or a *C. elegans* native protein library (right) were followed by MS-based phosphoproteomics analyses. The activity of a constitutively active mutant (CMK-1(1-295)T179D) was compared to control situations with either kinase dead mutant (CMK-1(1-295)K52A) or without ATP co-substrate during the incubation. **B**. Comparison of the results with the two approaches, showing the number of overlapping and non-overlapping phosphosites, the number of corresponding proteins, as well as the 15-residue consensus sequence surrounding the phosphosites. Logos created with Seq2Logo. **C**. Results of a separate *in vitro* kinase assay in which purified TAX-6/CnA was used as substrate and showing TAX-6/CnA S443 phosphorylation. N.D., not detected **D**. Diagram presenting the different regions of the *C. elegans* TAX6/CnA protein and the localization of S443. **E**. Alignments showing high conservation of a CnA protein region around S443 across *C. elegans, Danio rerio, Mus Musculus* and *Homo sapiens* (Uniprot assessions: Q0G819-2|PP2B_CAEEL; A3KGZ6|A3KGZ6_DANRE; P63328|PP2BA_MOUSE; Q08209|PP2BA_HUMAN).

With the peptide library data, we identified 646 phosphosites in 478 proteins (File S1). With the whole protein library, we identified 427 phosphosites in 365 proteins (File S1). Fifty-one phosphosites in 50 proteins were common between the two datasets (File S1, Fig.1B). As more extensively discussed later in the text (see Discussion section), these datasets are expected to include both phosphosites that are relevant CMK-1 target *in vivo* and ‘false positive’ phosphosites which are not relevant targets *in vivo.* These datasets are also expected to be biased toward abundant proteins, which are more likely to be detected. Consistent with this expectation, the most significant enrichment found via gene ontology (GO) term analyses in related to ribosomal and cytoskeletal proteins (File S2). Outside of these categories and among the phosphosites common to our two datasets, we were intrigued by the presence of a CMK-1 target phosphosite on the serine 443 of TAX-6/CnA. This phosphosite is located in a highly conserved region of TAX-6/CnA regulatory domain, just on the C-terminal side of the CaM binding domain (Fig. 1D). To confirm that CMK-1 can phosphorylate TAX-6/CnA S443 *in vitro*, we repeated the kinase assay using recombinant TAX-6/CnA protein purified from *E. coli* as substrate. Abundant phosphorylation of S443 was observed upon treatment with the constitutively active CMK-1 form, whereas this phosphorylation remained undetected with kinase-dead control treatment (Fig. 1C). We conclude that CMK-1 can phosphorylate TAX-6/CnA on S443 *in vitro*.

### Thermo-nociceptive adaptation is impaired by CMK-1 down-regulation and by both up- and down-regulation of TAX-6/CnA

We previously reported that CMK-1 controls the adaptation to persistent noxious heat stimulations (Schild, Zbinden et al. 2014) or repeated stimulations (Lia and Glauser 2020). This adaptation effect consists in a progressive reduction of the noxious heat-evoked reversal response observed in wild type during one hour of repeated heat stimulation. Importantly, this adaptation effect does not result from thermal damages or from an exhaustion of neuronal or muscular tissues, as evidenced by the existence of non-adapting mutants which can maintain a constant response level (Jordan and Glauser 2023). We confirmed these previous observations by quantifying heat-evoked reversals in naïve animals (T0) and animals submitted to a series of repeated heat stimulation for one hour (T60, *adaptation treatment* with an interstimulus interval (ISI) of 20 s, Fig. 2A). Whereas wild type response is significantly decreased following repeated stimulation treatment (1h adaptation, T60), this adaptation effect is absent in a *cmk-1(ok287)* loss-of-function mutants (Fig. 2B). Conversely, a *cmk-1(syb1633)* gain-of-function mutant with a CMK-1(T179D) over-activating mutation promoted a faster adaptation (Fig. 2B and Fig. 2-supplement 1). Therefore, our data confirm that CMK-1 activity is necessary and sufficient to promote thermo-nociceptive adaptation.

**Figure 2.**
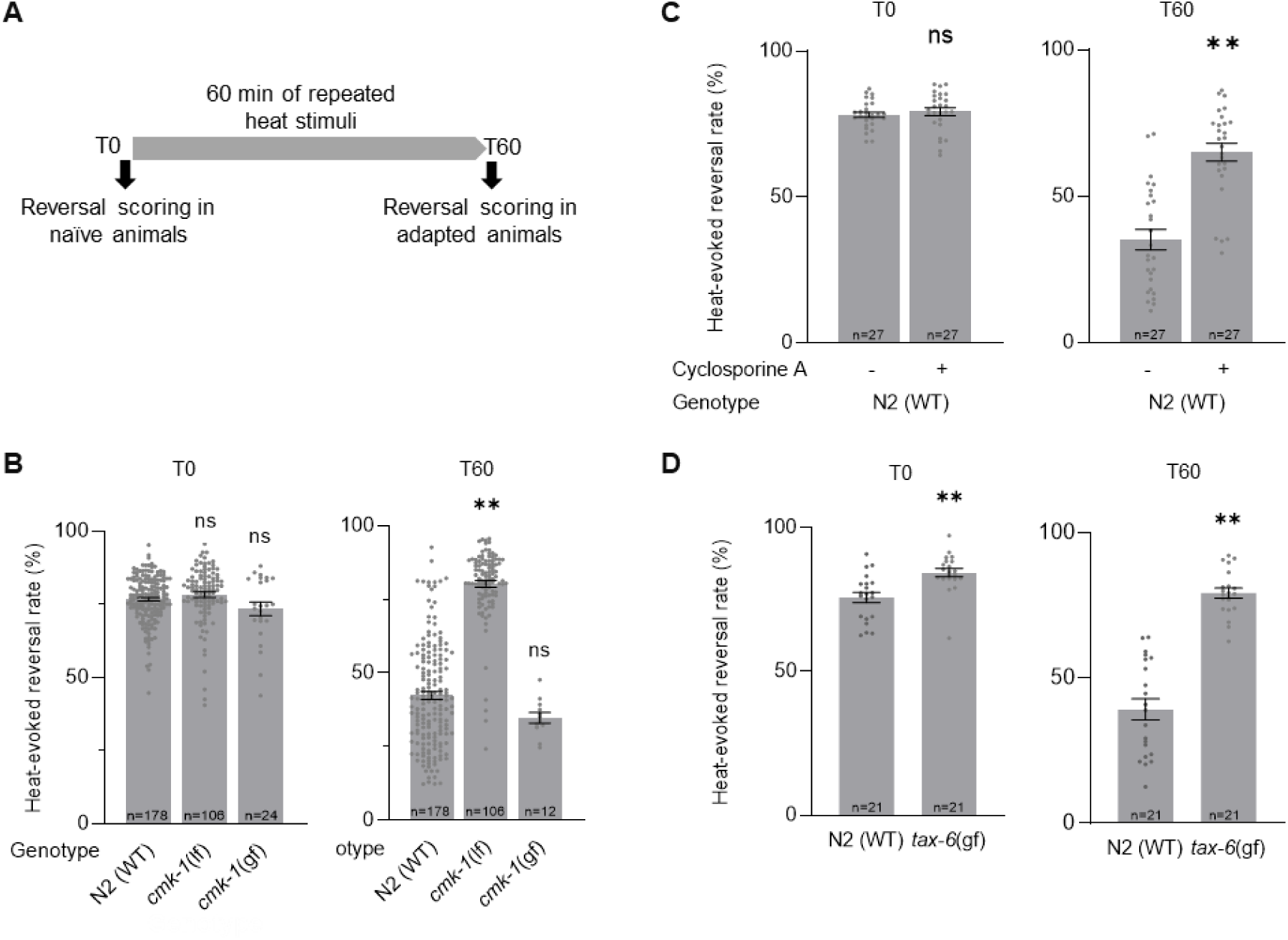
Impact of loss and gain of CMK-1 and TAX-6/CnA function on *C. elegans* thermo-nociceptive response. **A**. Schematic of the scoring procedure. Heat-evoked reversals were first scored in naïve adult *C. elegans* that had never been exposed to thermal stimuli (T0), animals exposed to 4 s heat pulses every 20 s during 60 min, prior to an endpoint scoring after adaptation (T60). **B-D**. Heat-evoked reversal scored in the indicated genotypes. Results as fraction of reversing animals. Each point corresponds to one assay scoring at least 50 animals. Average (grey bars) and s.e.m (error bars) with indicated *n* representing the number of independent assays. *cmk-1(gf)* is *cmk-1(syb1633)*. *cmk-1(lf)* is *cmk-1(ok287)*. *tax-6(gf)* is *tax-(j107)*. For Calcineurin inhibition, 10 µM Cyclosporin A was used 24 prior to experiments. **, *p*<.01 and *, *p*<.05 versus N2(WT) control in the specific condition by Bonferroni-Holm post-hocs tests. ns, not significant.

Since TAX-6/CnA is a CMK-1 kinase substrate *in vitro*, we hypothesized that it could take part in the regulation of thermo-nociceptive adaptation and evaluated the impact of genetic and pharmacological manipulations down-regulating or up-regulating calcineurin signaling activity. The permanent loss of TAX-6/CnA in *tax-6(p675)* mutants or of CnB/CNB-1 in a *cnb-1(jh103)* and *cnb-1(ok276)* mutants caused a marked elevation in the rate of spontaneous reversals (Fig. 2-supplement 2). While this observation indicates that TAX-6/CnA signaling regulates reversal behavior, the elevation was so high that it precluded quantifying heat-evoked reversals and plasticity response. We next turned to a pharmacological approach using the calcineurin inhibitor Cyclosporin A (Liu, Farmer et al. 1991). Treating wild type worms with 10 mM Cyclosporin A for 24 h prior to behavioral assays did not cause the problematic elevated spontaneous reversal phenotype, but significantly reduced the thermal adaptation effect (Fig.2C). An intact calcineurin signaling is therefore required for thermo-nociceptive adaptation. Next, we examined the impact of over-activating TAX-6/CnA, using a *tax-6(jh107)* gain-of-function mutant lacking the auto-inhibitory C-terminal domain of TAX-6/CnA, which we will refer to as *tax-6(gf)*. We noted a very slightly enhanced reversal response in naïve *tax-6(gf)* mutants in comparison to wild type (Fig. 2D). More strikingly, the adaptation effect in *tax-6(gf)* mutants was strongly impaired. Therefore, our data show that both overactivation and inhibition of TAX-6/CnA signaling can block thermo-nociceptive adaptation.

In summary, gain- and loss-of-function analyses highlight a CMK-1-dependent pathway promoting and potentially two antagonistic TAX-6/CnA-dependent pathways that promote and inhibit thermo-nociceptive adaptation, respectively.

### CMK-1 and TAX-6/CnA signaling regulates thermo-nociceptive adaptation through a set of inhibitory cross-talks

To assess the potential interactions between CMK-1 and calcineurin signaling in the control of thermo-nociceptive adaptation, we systematically tested combinations of up- or down-regulating manipulations in the two pathways.

First, we examined the impact of down-regulating both CMK-1 and TAX-6/CnA signaling by treating *cmk-1(lf)* mutants with Cyclosporin A. Surprisingly, whereas separate down-regulation of either CMK-1 or TAX-6/CnA pathway strongly blocked adaptation, their joint down-regulation caused animals to adapt like wild type (Fig. 3A). An intact adaptation when both pathways are inhibited supports the notion that CMK-1 and TAX-6/CnA represent regulators of one or more adaptation pathways that can also operate independently. In addition, this observation supports a model in which (i) the anti-adaptation effect caused by CMK-1 inhibition is mediated by TAX-6/CnA and, conversely, (ii) that the anti-adaptation effect caused by TAX-6/CnA inhibition requires intact CMK-1 activity.

**Figure 3.**
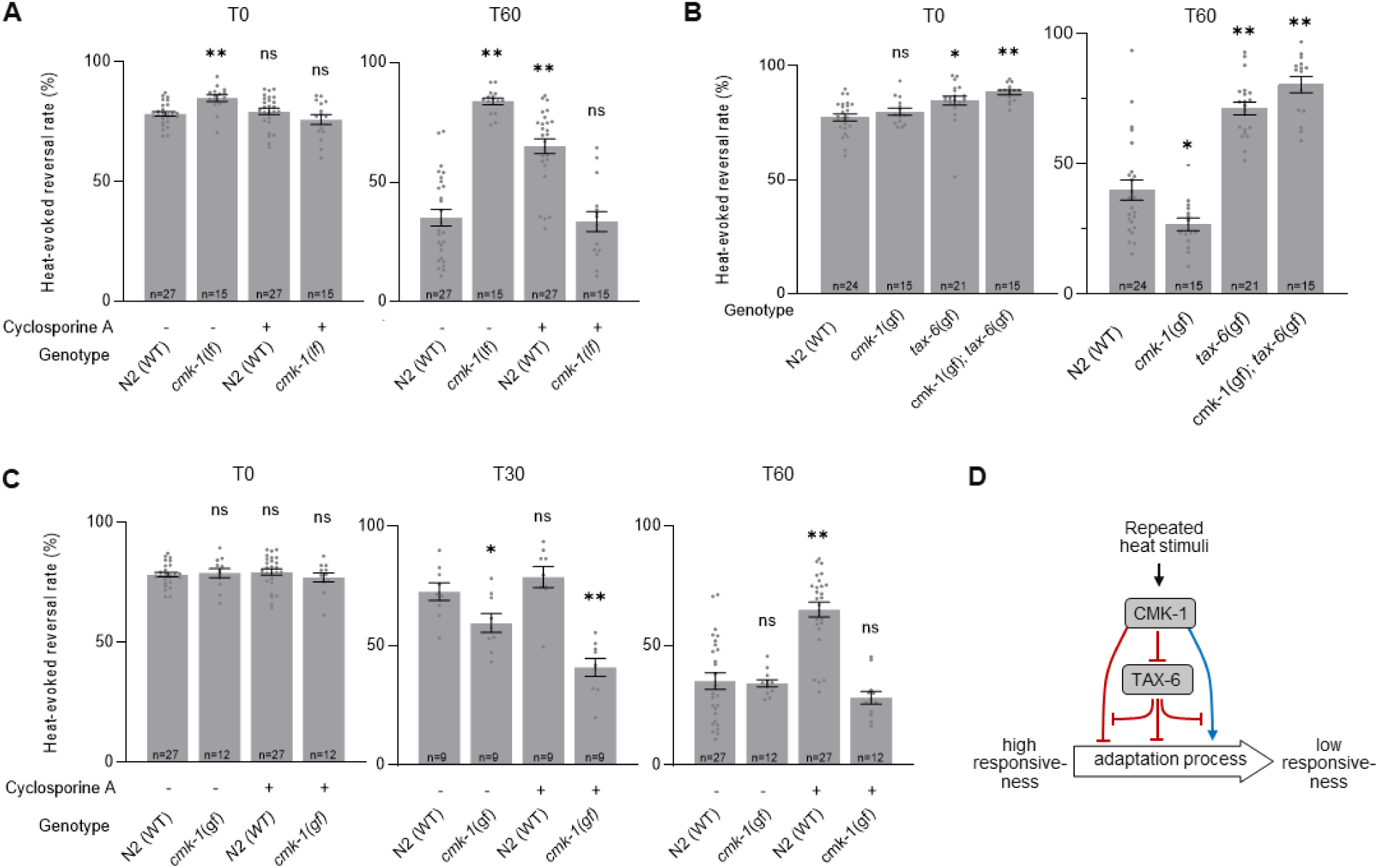
Functional interactions between CMK-1 and TAX-6/CnA in the regulation of thermo-nociceptive adaptation. **A-C**. Assessment of the impact of joint gain- and loss-of-function manipulations affecting the CMK-1 and TAX-6/CnA pathways. Heat-evoked response in naïve animals (T0) and after 60 min of repeated stimulations (T60), scored and reported as in Fig. 2. **, *p*<.01 and *, *p*<.05 versus N2(WT) control in the specific condition by Bonferroni-Holm post-hocs tests. ns, not significant. **D**. Schematic of a model explaining the multiple antagonistic interactions observed between CMK-1 and TAX-6/CnA signaling. *cmk-1(gf)* is *cmk-1(syb1633)* harboring the CMK-1(T179D) overactivating mutation. *cmk-1(lf)* is *cmk-1(ok287)*. *tax-6(gf)* is *tax-6(jh107)*.

Second, we tested the impact of simultaneously activating CMK-1 and calcineurin pathways by examining the behavior of *cmk-1(gf);tax-6(gf)* double mutants. We found that the double mutants behaved like *tax-6(gf)* single, entirely lacking adaptation (Fig. 3B). Therefore, the pro-adaptation effect of the CMK-1(T179D) activating mutation is fully blocked by TAX-6/CnA overactivation, which is compatible with a model in which CMK-1 over-activation might work by inhibiting calcineurin signaling.

Third, we assessed the impact of concomitantly up-regulating calcineurin and down-regulating CMK-1 signaling, by examining the behavior of *cmk-1(lf);tax-6(gf)* double mutants. Whereas the main phenotypic feature in each single mutant was a lack of adaptation, the mutation combination produced a synthetic effect massively elevating spontaneous reversal, which precluded quantifying heat-evoked reversals (Fig. 2-supplement 2).

Fourth, we tested the effect of concomitantly up-regulating CMK-1 activity and inhibiting calcineurin signaling by treating *cmk-1(gf)* mutants with Cyclosporin A. We found that, unlike in *cmk-1(wt)* background, Cyclosporin A treatment in *cmk-1(gf)* mutants did not prevent thermo-nociceptive adaptation (Fig.3C). The fact that an overactive CMK-1 signaling can compensate for the inhibition of TAX-6/CnA suggests that the anti-adaptation effect of calcineurin inhibition involves the ability to down-regulate CMK-1 signaling. Furthermore, the ability of CMK-1 overactivation to promote adaptation when calcineurin is inhibited indicate that at least one CMK-1 regulatory branch works independently or downstream of calcineurin signaling.

Collectively, our results do not support a simple linear model in which CMK-1 would only work upstream of TAX-6/CnA. Instead, our data suggest a model in which thermo-nociceptive adaptation processed are regulated via the antagonist actions of CMK-1 and TAX-6/CnA signaling operating through a non-linear inhibitory network, such as the one depicted in Figure 3D and discussed below (see Discussion section).

### CMK-1 acts in AFD and ASER sensory neurons to promote thermo-nociceptive adaptation

Our next goal was to assess the place of action of CMK-1 signaling in the promotion of thermo-nociceptive adaptation. To that end we used a neuron-specific rescue approach in the *cmk-1(lf)* background. We started our experiments by restoring CMK-1 expression in candidate thermo-sensory neurons, using *mec-3p* promoter to drive expression in FLP and *tax-4p* promoter to drive expression in AFD, AWC, ASI and ASE, as well as *glr-1p* promoter to drive expression in a subset of interneurons known to mediate reversal response. A positive control using *cmk-1p* promoter showed a partial, yet significant rescue effect (Fig. 4A). We observed a strong rescue effect with the *tax-4p* promoter, but no rescue effect with *mec-3p* or *glr-1p* (Fig. 4A). While not ruling out a contribution of other neurons, these data point to *tax-4-*expressing neurons as a major place of action for CMK-1 in the control of thermo-nociceptive adaptation.

**Figure 4.**
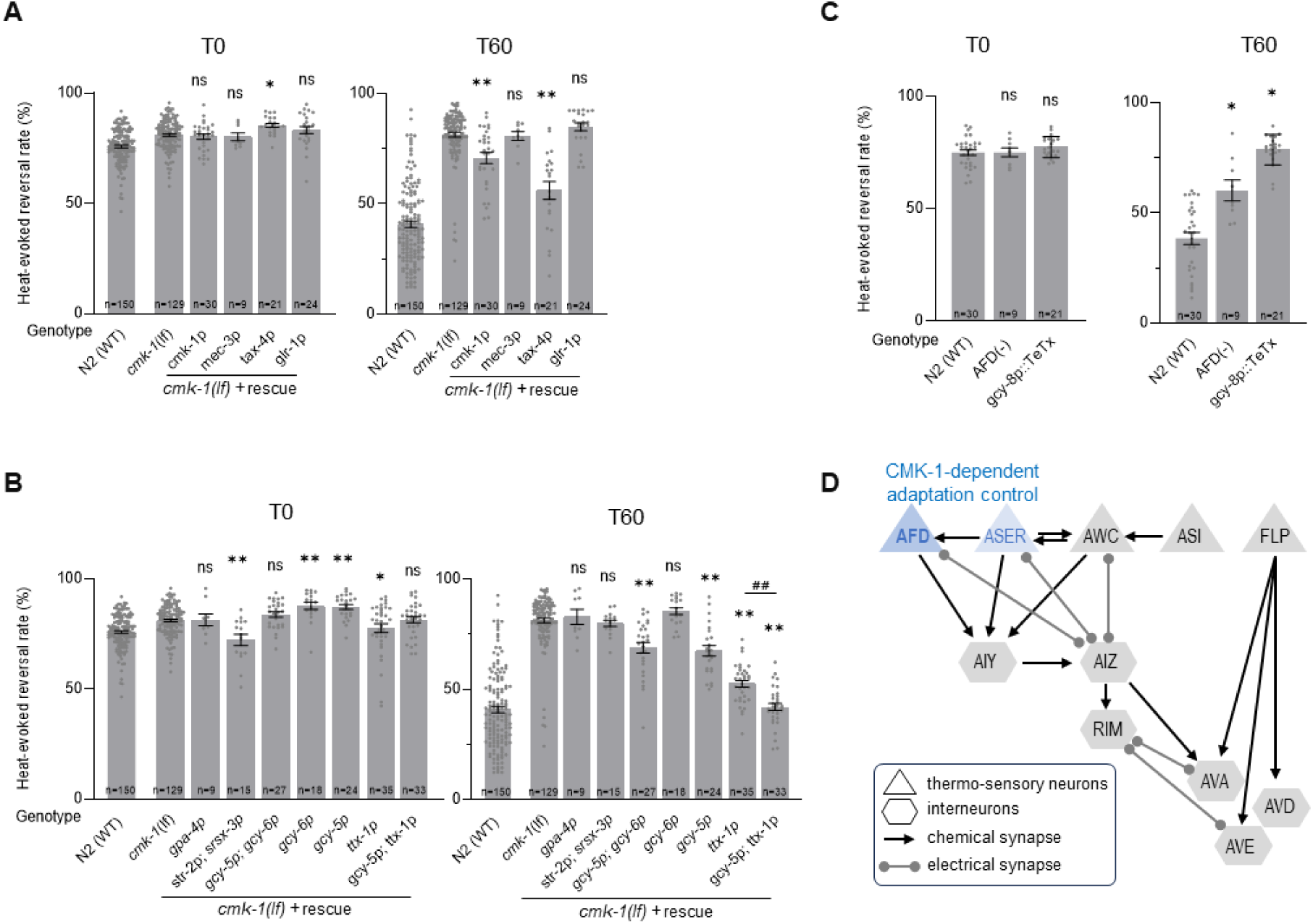
CMK-1 works in AFD and ASER to control thermo-nociceptive adaptation. **A-B**. Determination of the CMK-1 place of action in the control of thermo-nociceptive adaptation, using cell-specific rescue in *cmk-1(ok287)* (*cmk-1(lf)*) background. Heat-evoked response in naïve animals (T0) and after 60 min of repeated stimulations (T60), scored and reported as in Fig. 2. The promoters used to restore CMK-1 expression are indicated below each bar. **, *p*<.01 and *, *p*<.05 versus *cmk-1(lf)* by Bonferroni-Holm post-hocs tests. ns, not significant. ##, *p<*.01 for the specific contrast between *ttx-1p* and the *gcy-5p;ttx-1p* combination. N2(WT) data are shown for comparison purpose. **C**. Impact of genetic manipulation ablating AFD with a caspase construct (AFD(-)) or inhibiting AFD neurotransmission with TeTx heterologous expression. **, *p*<.01 and *, *p*<.05 versus N2(WT) control by Bonferroni-Holm post-hocs tests **D**. Schematic of the hypothetical circuit controlling noxious-heat evoked reversals, including the *tax-4*-expressing thermo-responsive sensory neurons AFD, AWC, ASER and ASI, the *mec-3*-expressing FLP thermo-nociceptor, and a subset of downstream interneurons known to mediate reversal response, including the *glr-1-*expressing RIM, AVA, AVD, and AVE interneurons. The main loci of action of CMK-1 determined through the cell-specific rescue approach are highlighted in blue.

To further dissect the contribution of *tax-4-*expressing neurons, we conducted additional rescue experiments using *gpa-4p* to target ASI, a combination of *str-2p* and *srsx-3p* to target the two AWC neurons (AWCon and AWCoff, respectively), a combination of *gcy-5p* and *gcy-6p* to target ASEL and ASER neurons, *gcy-6* alone to target ASEL, *gcy-5p* alone to target ASER and *ttx-1p* to target AFD. We observed an almost total rescue effect with the *ttx-1p* promoter (AFD) and a more partial rescue effect with either *gcy-5p+gcy-6p* (both ASEL and ASER) or *gcy-5p* alone (only ASER) (Fig. 4B). No rescue effect was observed with the other promoters. We additionally tested the combined use of *ttx-1p* and *gcy-5p* to target both AFD and ASER and obtained a maximal rescue effect even stronger than with *ttx-1p* alone (Fig. 4B). Taken together, out data point to AFD as a major (and to ASER as a more minor) place of action for CMK-1 signaling in the control of thermo-nociceptive adaptation to repeated heat-stimuli.

To further asses the role of AFD neurons, we evaluated the behavior of animals with genetically ablated AFD neurons. Interestingly, AFD neurons were dispensable for heat-evoked reversal responses in naïve animals under our experimental conditions, but required for the thermo-nociceptive adaptation (Fig. 4.C). We observed similar results when selectively inhibiting AFD neurotransmission in animals expressing the tetanus toxin (TeTx) in AFD (Fig. 4C).

Taken together, our results show that AFD neurons mediate thermo-nociceptive adaptation and that intact CMK-1 signaling and neurotransmission in these neurons is essential for this process (Fig. 4D). ASER might represent an additional locus of CMK-1 action with a milder quantitative contribution to the process (Fig. 4D).

### TAX-6/CnA activity in RIM and command interneurons inhibits thermo-nociceptive adaptation

In order to assess the cellular place of action of TAX-6/CnA in the control of thermo-nociceptive adaptation, we used a neuron-type-specific overactivation approach. We created transgenes containing a *tax-6(gf)* cDNA to drive the expression of the overactive, truncated form of TAX-6/CnA encoded in the *tax-6(jh107)* mutant. Like for CMK-1 rescue experiments above, we started by targeting candidate thermosensory neurons using either the *mec-3p* (FLP) or *tax-4p* (AFD, AWC, ASI and ASE) promoters, as well as interneurons with the *glr-1p* promoter. We examined the ability of the transgenes to replicate the two noticeable phenotypes of *tax-6(gf)* mutants: a slightly enhanced response in naïve animals and a strong impairment of adaptation.

First, regarding naïve animal response, we found that *[mec-3p::tax-6(gf)]* and, to a lesser extent *[glr-1p::tax-6(gf)],* transgene caused a slight elevation in the heat-evoked response, whereas *[tax-4p:tax-6(gf)]* did not produce any effect (Fig. 5A). *glr-1p* promoter drives expression in several interneurons, including AVA, AVD, AVE and RIM whose activity is required and sufficient to trigger reversals (Alkema, Hunter-Ensor et al. 2005, Gray, Hill et al. 2005, Guo, Hart et al. 2009). To further dissect the specific interneurons involved, we expressed the *tax-6(gf)* transgene in AVA, AVD and AVE using the *nmr-1p* promoter, or the *lgc-39p* promoter, in RIM (and RIC) using the *cex-1p* promoter, and only in RIM using the *tdc-1p* promoter (Brockie, Mellem et al. 2001, Lemieux, Cunningham et al. 2015, Thapliyal, Beets et al. 2023). With all four promoters, we observed an effect similar to that with *glr-1p*. Taken together, these results indicate that over-activating calcineurin signaling in *mec-3p*-expressing neuron (most likely in FLP thermo-nociceptor) or in a subset of reversal-mediating interneurons such as RIM or AVA/AVD/AVE is sufficient to up-regulate noxious heat-responsiveness in naïve animals (Fig. 5C, top).

**Figure 5.**
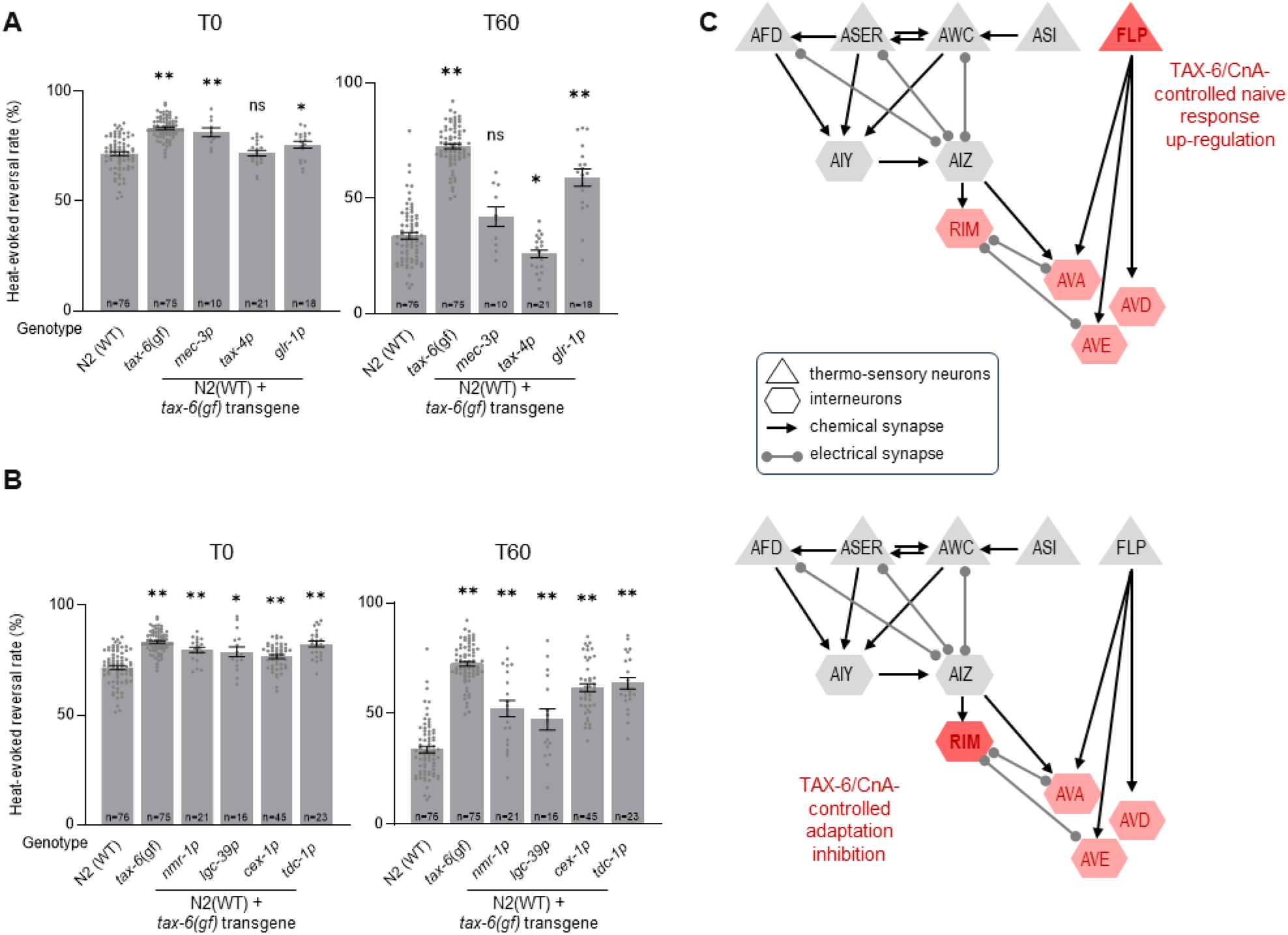
TAX-6/CnA activity in RIM and AVA/AVD/AVE inhibit thermo-nociceptive adaptation. **A-B**. Determination of the TAX-6/CnA place of action in the control of thermo-nociceptive responses, using cell-specific expression of a TAX-6/CnA gain-of-function mutant. Heat-evoked response in naïve animals (T0) and after 60 min of repeated stimulations (T60), scored and reported as in Fig. 2. The promoters used to drive *tax-6(gf)* cDNA expression are indicated below each bar. **, *p*<.01 and *, *p*<.05 versus N2(WT) by Bonferroni-Holm post-hocs tests. ns, not significant. *tax-6(gf)* mutant data are shown for comparison purpose. **C**. Schematics of the hypothetical circuit controlling noxious-heat evoked reversals as in Fig. 4D, with the main loci of action of TAX-6/CnA highlighted in red. TAX-6/CnA-evoked up-regulation of thermo-nociceptive response in naïve animals (top) and TAX-6/CnA-evoked inhibition of adaptation (bottom).

Second, regarding adaptation to repeated heat stimuli, we found that the *[glr-1p::tax-6(gf)]* transgene could significantly reduce the adaptation effect, leaving stronger response as compared to wild type after one hour of repeated stimulation (Fig. 5A). This effect was not as strong as the effect in *tax-6(gf)* mutant, which could be due to the transgene overexpression, mosaicism in the transgenic animals or to the fact that TAX-6/CnA overactivation in additional (unidentified) neurons could be also needed to reach the full effect. In contrast, the *[mec-3p::tax-6(gf)]* transgene did not affect nociceptive adaptation, whereas the *[tax-4p::tax-6(gf)]* transgene produced a slight enhancement of adaptation (p<.05). The data in Fig. 5A suggest that one or more *glr-1-*expressing interneurons represent a major place of action in which overactive TAX-6/CnA signaling can act to inhibit adaptation. We deepened the circuit analysis with additional promoters and observed a marked adaptation defect caused with *cex-1p* and *tdc-1p* promoters, similar to that with *glr-1p* promoter, and a milder defect with *lgc-39p* and *nmr-1p* promoter. These results point to RIM second layer interneurons as primary locus and AVA/AVD/AVE reversal command neurons as secondary locus of overactive TAX-6/CnA action, while not ruling out the implication of additional neurons (Fig. 5C bottom).

## DISCUSSION

### CMK-1 substrate specificity is indistinguishable from that of mammalian CaMKs

Our knowledge on CaMK substrate specificity comes from numerous previous biochemical studies on mammalian CaMKs using synthetic peptide substrates at different scales (Lee, Kwon et al. 1994, White, Kwon et al. 1998, Corcoran, Joseph et al. 2003, Johnson, Yaron et al. 2023). Our study complements and extends this previous work through a large-scale identification of CaMKI substrates with endogenous peptide and protein libraries. With a dataset comprising several hundred substrates, we could confirm the ΦXRXX(S/T)XXXΦ consensus (where Φ represents hydrophobic residues) (Lee, Kwon et al. 1994, White, Kwon et al. 1998). These findings suggest that (i) the vast majority of our hits are direct CMK-1 substrates *in vitro* and (ii) the substrate specificity is conserved from worm to mammals.

### Large-scale empirical identification of CMK-1 phospho-targets

We have identified several hundred substrates that can be directly phosphorylated by CMK-1 *in vitro*. Our hit list is expected to include both substrates that are relevant *in vivo* and substrates that are not. Indeed, both the peptide library and the whole protein library datasets could include phosphosites coming from proteins that never encounter CMK-1 *in vivo*: e.g., non-neuronal proteins, secreted proteins, or proteins residing in specific organelles. Furthermore, both datasets are expected to be biased toward abundant proteins, which are more likely to be detected. Accordingly, the strongest GO term enrichments in both datasets related to abundant and ubiquitous cellular components, involved in the translation process and cytoskeleton. Even if we cannot rule out an actual inclination of the CaM kinase pathway to regulate these processes, we suspect that these GO term enrichments rather reflect an analytical bias toward abundant proteins.

The relatively low overlap between the peptide and the protein library datasets can be explained by multiple effects. First, the peptide library dataset is expected to include phosphosites in protein regions that are usually inaccessible to CMK-1 because of protein folding or protein complex formation (increasing the false positive rate in the peptide library). Second, the trypsin-bases cleavage prior to CMK-1 exposure in the peptide library protocol might impair proper substrate recognition by the kinase (increasing false negative rate in the peptide library). Third, we have chosen quite strict criteria and applied them separately to define each hit list. For theoretical reasons, because substrates were presented to the kinase in a more native condition, we tend to give more trust the protein library dataset, with the subset of targets also found in the peptide library as top hits.

A previous study reported a list of 533 CMK-1 candidate substrates based on *in silico* predictions (Ardiel, McDiarmid et al. 2018). We found very little overlap between those predictions and our empirical datasets, with only 5 phosphosites shared with our peptide library list and 1 with our protein library list (Fig. 1-supplement 1). This limited overlap could be explained either by a limited exhaustiveness across the three datasets (high false negative rate) or come from limitations affecting *in silico* predictions. In support of the second explanation, we note that the previously predicted phosphosite list only very partially matches the empirically determined CMK-1 substrate consensus (Fig. 1-supplement 1). Because we lack a solid reference point, it is nevertheless hard to estimate the false positive and false negative rates in our analyses at this stage. Additional experiments will be required to determine if the phosphorylation of these substrates is relevant *in vivo*, potentially on a case-by-case basis. Eventually, and keeping these limitations in mind, we believe that our large-scale dataset on the substrates which can by phosphorylated by CMK-1 *in vitro* will represent a useful resource in the search for CMK-1/CaMKI substrates mediating its numerous biological functions *in vivo*.

### The phosphorylation of TAX-6/CnA by CMK-1/CaMKI

Previous *in vitro* studies have found that CnA can be phosphorylated by protein kinase C, casein kinase I, casein kinase II, protein kinase C and by CaM kinase II (Hashimoto, King et al. 1988, Hashimoto and Soderling 1989, Calalb, Kincaid et al. 1990, Rusnak and Mertz 2000). We are not aware of any study having shown that CnA is a substrate of CaM kinase I. Here, we show that *C. elegans* CaMKI CMK-1 can phosphorylate TAX-6/CnA on S443. This specific phosphorylation event was confirmed in three different types of kinase assays: one using a peptide library, one using a protein library, and one using purified recombinant TAX-6/CnA as substrate. Ser 443 is located just downstream of the CaM binding domain in a highly conserved region (Fig. 1D and E). This conserved serine was previously identified as a target of mammalian CaMKII *in vitro* (Hashimoto, King et al. 1988, Hashimoto and Soderling 1989). Because TAX-6/CnA and CMK-1 seem to mostly work from distinct neuronal cell-type to control nociceptive adaptation, it remains unclear whether TAX-6/CnA phosphorylation by CMK-1 is directly relevant for the control of this process *in vivo*. Nevertheless, since the expression patterns of CMK-1 and TAX-6/CnA largely overlap and since they are both activated by calcium signaling, we speculate that this phosphorylation event might be relevant to other neuronal regulatory signaling and we suggest it should be considered in future studies addressing the potential cross-talks between the two pathways.

### Complex interaction network and distributed cellular locus of actions for CMK-1 and Calcineurin signaling

Our systematic analysis of CMK-1 and calcineurin signaling cross-talks has revealed a relatively complex regulatory network in which the two pathways mostly antagonize each other to regulate adaptation. As mentioned above, adaptation still takes place when CMK-1 and TAX-6/CnA are concomitantly inhibited, which led us to model their action as regulatory events controlling a separate adaptation process (see model in Fig. 3D). While relatively complex, this model is the simplest we could articulate to explain all the empirically measured interactions (see Fig.3-supplement 1 for a case-by-case illustration of the proposed regulatory network functioning in our different experimental conditions). In this model network, CMK-1 can regulate thermal nociception via three pathways: in the first pathway CMK-1 promotes adaptation by inhibiting TAX-6/CnA, which in turn inhibits adaptation. In the second and third pathways, CMK-1 works independently of TAX-6/ CnA to inhibit and promote adaptation, respectively. These two latter pathways can each be gated by TAX-6/ CnA. Therefore, both CMK-1 and calcineurin signaling activities may promote or inhibit adaptation and shifting their activity balance could represent a way to achieve nuanced yet robust modulation, integrating past activity with potentially additional cues. This complex regulation scheme would be hard to achieve if the two signaling pathways were working in a single neuronal cell type and it was therefore not surprising that our cell-specific approaches revealed multiple cellular loci of action in the circuit controlling temperature sensation and reversal execution.

In a previous study, CMK-1 was shown to work cell-autonomously in FLP to modulate thermo-nociceptive responses following persistent noxious heat stimulation (Schild, Zbinden et al. 2014). FLPs are ‘tonic’ thermo-sensory neurons, whose activity constantly reflects the current temperature (Saro, Lia et al. 2020). Here, we show that AFDs, but not FLPs, constitute the main cellular locus of action of CMK-1-dependent plasticity in the case of repeated short-lasting stimulations. Interestingly, AFD are mostly ‘phasic’ thermo-sensory neurons producing activity peaks in response to thermal changes (Kimura, Miyawaki et al. 2004, Clark, Biron et al. 2006, Hawk, Calvo et al. 2018, Glauser 2022). They are thus ideally suited to encode repeated thermal pulses and modulate the heat-evoked reversal circuit. Laser-ablation of AFD was previously shown to reduce reversal in response to head-targeted infra-red laser beams (Liu, Schulze et al. 2012). Here, we used thermal stimuli that were diffuse (whole animal exposure) and found that neither the genetic ablation of AFD, nor the cell-specific inhibition of neurotransmission affected the animal’s ability to produce heat-evoked reversals. Instead, AFD was essential to favor an experience-dependent reduction in heat-evoked reversal response. CMK-1 was previously shown to mediate both short-term (minute timescale) and long-term (hour timescale) adaptation in AFD intracellular calcium activity that resulted from thermal changes within the 15-25 °C innocuous temperature range (Yu, Bell et al. 2014). Long-term changes involved gene expression changes, whereas the processes mediating short-term adaptation remained elusive (Yu, Bell et al. 2014). At this stage, we do not know exactly how CMK-1 works in AFD to orchestrate thermo-nociceptive adaptation, but we can envision many possible mechanisms, including qualitative or quantitative modulation of AFD thermo-sensitivity, neuronal excitability, calcium dynamics, neurotransmitter or neuro-modulator release, or even developmental effect affecting AFD synaptic connectivity. Likewise, we don’t know the downstream circuit involved. Among different possibilities the CMK-1 signaling taking place in AFD could affect reversal via AFD’s direct interneuron partners AIZ and/or AIY, or engage extra-synaptic communication with neuropeptides. Additional studies will thus be needed to address the downstream processes that are engaged within and downstream of AFD to modulate animal responsiveness to noxious heat stimuli.

We found that calcineurin signaling works at multiple cellular loci to control thermo-nociceptive response. On the one hand, overactive TAX-6 activity in FLP is sufficient to increase reversal response in naïve animals. Since FLP activity is known to favor reversal response, we might hypothesize that calcineurin activity could promote FLP activity. On the other hand, overactive TAX-6 activity in RIM or, to a lesser extent, in AVA/AVD/AVE neuron is sufficient to increase reversal response in naïve animals, but conversely enhances reversal response upon repeated stimulation. This suggests that the impact that calcineurin signaling has in these neurons depends on past experience. AVA/AVD/AVE are thought to mostly promote reversal response, whereas RIM neuron activity might either up-regulate, or down-regulate reversal responses based on the context (Alkema, Hunter-Ensor et al. 2005, Gray, Hill et al. 2005, Guo, Hart et al. 2009, Piggott, Liu et al. 2011, Cho and Sternberg 2014, Li, Zhou et al. 2023). One particularly interesting observation is that cell-autonomous TAX-6 overactivation in interneurons acts as a major gate blocking adaptation irrespective of any CMK-1 activity manipulation. One hypothetical mechanism could be that TAX-6/CnA activity in worm would work in post-synaptic region of interneuron to adjust synaptic strength, e.g., via its action on ion channels or on inhibitory or excitatory neurotransmitter receptors, as shown in various models of synaptic plasticity in mammals (Groth, Dunbar et al. 2003).

## Conclusion

In summary, our study reports the empirical identification of many potential CMK-1 targets *in vitro*, among which the catalytic subunit of Calcineurin TAX-6/CnA. Whereas we have not yet found evidence that the phosphorylation of TAX-6 by CMK-1 is directly relevant for thermo-nociceptive response modulation, we show a complex interplay between CMK-1 and calcineurin signaling that operate via multiple regulatory nodes within the noxious-heat-evoked reversal circuits of *C. elegans*. Our study paves the role for a deeper dissection of how conserved intracellular signaling pathways operating at distributed loci within a sensory-behavior circuit can actuate experience-dependent changes in nociceptive behavior.

## MATERIAL AND METHODS

### Worm maintenance

All *C. elegans* strains used in this study are listed in File S4. All strains were grown on nematode growth media (NGM) plates with OP50 *E. coli* (Stiernagle, 2006) at 20°C. For TAX-6 inhibition experiments, NGM plates containing cyclosporine A were used, as well as regular NGM plates as control. Cyclosporine A (10 μM) plates were prepared 72h before experiments and kept protected from light at room temperature.

### Expression and purification of CMK-1(T179D)-GST, CMK-1(K52A)-GST and TAX-6-HIS6

DNA fragments encoding CMK-1 and TAX-6 proteins were PCR amplified and cloned into NdeI and BamHI restriction sites of pDK2409 (pET-24d/GST-TEV-KAP104.419C) for the GST-TEV-tagged protein and of pDK2832 (pET-24d/(His)6) for the His6-tagged proteins. Plasmids were transformed into *E. coli* BL21 (DE3) (Novagen). For protein expression, bacteria were grown overnight at 37°C, next day transferred to 200 ml of Luria Broth (LB) with kanamycin 30 μg/ml to OD_600_=0.1, incubated up to OD_600_=0.5. Then, protein expression was induced using 0.5 mM isopropyl β-D-1-thiogalactopyranoside (IPTG) and incubated for 5 h at 23 °C. After, bacteria cells were collected by centrifugation at 2800 rcf for 10 min at 4°C, resuspended in cold lysis buffer (150 mM NaCl, 50 mM Tris-HCl pH7.5, 1.5 mM MgCl_2_, 5% Glycerol, 1 mM phenylmethylsulfonyl fluoride (PMSF)) and lysed in a Microfluidizer Processor M-110L. Then, NP-40 was added to the concentration of 0.1%, and soluble extract was obtained by centrifugation at 23’400 rcf for 20 min at 4 °C. Supernatant was incubated for 2 h at 4°C while rotating with Glutathione superflow beads (Qiagen) for GST-tagged CMK-1 variants or nickel-nitrilotriacetic acid beads (Ni-NTA, Qiagen) in the presence of 15 mM imidazole for TAX-6-His6 protein binding. Beads were harvested at 580 rcf at 4 °C. GST-tagged CMK-1 kinase was cleaved from the tag during incubation in lysis buffer (added 1 mM PMSF, 0.1% NP-40, 1 mM Dithiothreitol (DTT)) with home-made TEV protease. TAX-6-bound Ni-NTA beads were washed 7 times with imidazole-containing buffer (5x with 15 mM and 2x with 50 mM imidazole) and eluted in lysis buffer with 1 mM PMSF, 0.1% NP-40, 500 mM imidazole. Protein concentration was determined by Pierce Microplate BCA protein assay Kit-Reducing Agent Compatible (Thermo Scientific) using BSA as protein standard.

### Total protein lysate preparation

Worms were grown on 10 cm diameter plates to the stage of young adults, washed with distilled water and suspended in extraction buffer. For *in vitro* kinase assays on peptide, urea buffer (8 M urea, 50 mM Tris-HCl pH 8.5) was used. For *in vitro* kinase assays on intact proteins, native protein extraction buffer (50 mM HEPES pH 7.4, 1% NP-40, 150 mM NaCl, 1x protease inhibitors, 0.5 mM PMSF) was used. Worms were flash-frozen in liquid nitrogen and cryogenically disrupted by using a Precellys homogenizer and acid-washed glass beads (5000 x 3 for 30 s with 30 s pause after each cycle). Lysates were collected by centrifugation at 1500 rcf at 4 °C.

### *In vitro* kinase assay

Purified CMK-1 kinase was used the same day to avoid freezing. *In vitro* kinase assays were performed according to (Hu, Sankar et al. 2021). Briefly, for kinase assays performed on native proteins, 30 mg of worm protein extract was incubated with pre-washed NHS-activated Sepharose beads at 4°C on a rotor for 6 h. Beads were washed with 3 × 10 ml of phosphatase buffer (50 mM HEPES, 100 mM NaCl, 0.1% NP-40). 1 ml of phosphatase buffer containing 5,000 to 10,000 units of lambda phosphatase with 1 mM MnCl_2_ was added and incubated for 4 h at room temperature (RT) followed by overnight incubation at 4°C on the rotor to dephosphorylate endogenous proteins. Beads were washed with 2 × 10 ml of kinase buffer (50 mM Tris-HCl pH 7.6, 10 mM MgCl_2_, 150 mM NaCl and 1x PhosSTOP (Roche)). Endogenous kinases bound to beads were inhibited by incubation with 1 mM FSBA in 1 ml of kinase buffer at RT on the rotor for 2 h. In addition, inhibition of the remaining active kinases was achieved with a further 1 h incubation in the presence of staurosporine (LC Laboratories), added to a final concentration of 100 μM. The beads were washed with 3 × 10 ml of kinase buffer to remove non-bound kinase inhibitors. The supernatant was removed completely using gel loading tips. Beads were split into 6 tubes (3x with kinase and 1 mM ATP, 3x with kinase and without ATP (Sigma-Aldrich), kinase buffer was added and incubated for 4 h at 30°C while shaking. For TAX-6 phosphorylation by CMK-1 *in vitro* assay, CMK-1 was kept on Glutathione superflow beads and purified TAX-6 protein in kinase buffer (50 mM HEPES pH7.5, 10 mM Mg(Ac)_2_, 1 mM DTT, 1x PhosSTOP) was added onto protein/bead mix together with ATP. Afterwards samples were lyophilized followed by protein digestion using trypsin. Samples were chemically labeled using dimethyl-labeling (Boersema, Raijmakers et al. 2009) supporting relative MS-based quantification. For kinase assays performed on peptide libraries, the procedure was the same except that protein extract samples were digested with Lyc-C (Lysyl Endopeptidase, WAKO) for 4 h and trypsin (Promega) overnight, before incubating them with Sepharose beads. Later beads were incubated either with purified active CMK-1 or kinase dead CMK-1 in kinase buffer (50 mM Tris-HCl pH 7.6, 10 mM MgCl_2_, 150 mM NaCl, 1x PhosSTOP, 1 mM DTT and 1 mM γ-[^18^O4]-ATP (Cambridge Isotope Laboratories)).

Peptides were fractionated and phosphopeptides enriched as described (Hu, Sankar et al. 2021); briefly: samples were incubated with TiO2 (GL Sciences) slurry, which was pre-incubated with 300 mg/ml lactic acid in 80% acetonitrile, 1% trifluoroacetic acid (TFA) prior to enrichment for 30 min at RT. For elution, TiO2 beads were transferred to 200 μl pipette tips blocked by C8 discs. After washing with 10% acetonitrile/1% TFA, 80% acetonitrile/1% TFA, and LC-MS grade water, phosphopeptides were eluted with 1.25% ammonia in 20% acetonitrile and 1.25% ammonia in 80% acetonitrile. Eluates were acidified with formic acid, concentrated by vacuum concentration, and resuspended in 0.1% formic acid for LC-MS/MS analysis.

#### LC-MS/MS analyses

LC-MS/MS measurements were performed on two LC-MS/MS systems as described (Hu, Sankar et al. 2021): a QExactive (QE) Plus and HF-X mass spectrometer coupled to an EasyLC 1000 and EasyLC 1200 nanoflow-HPLC, respectively (all Thermo Scientific). Peptides were fractionated on fused silica HPLC-column tips using a gradient of A (0.1% formic acid in water) and B (0.1% formic acid in 80% acetonitrile in water): 5%–30% B within 85 min with a flow rate of 250 nl/min. Mass spectrometers were operated in the data-dependent mode; after each MS scan (m/z = 370 – 1750; resolution: 70’000 for QE Plus and 120’000 for HF-X) a maximum of ten, or twelve MS/MS scans were performed (normalized collision energy of 25%), a target value of 1’000 (QE Plus)/5′000 (HF-X) and a resolution of 17’500 for QE Plus and 30’000 for HF-X. MS raw files were analyzed using MaxQuant (version 1.6.2.10) (Cox and Mann 2008) using a full-length *C. elegans* Uniprot database (November, 2017), and common contaminants such as keratins and enzymes used for in-gel digestion as reference. Carbamidomethylcysteine was set as fixed modification and protein amino-terminal acetylation, serine-, threonine- and tyrosine- (heavy) phosphorylation, and oxidation of methionine were set as variable modifications. In case of labeled samples, triple dimethyl-labeling was chosen as quantification method (Boersema, Raijmakers et al. 2009). The MS/MS tolerance was set to 20 ppm and three missed cleavages were allowed using trypsin/P as enzyme specificity. Peptide, site, and protein FDR based on a forward-reverse database were set to 0.01, minimum peptide length was set to 7, the minimum score for modified peptides was 40, and minimum number of peptides for identification of proteins was set to one, which must be unique. The “match-between-run” option was used with a time window of 0.7 min. MaxQuant results were analyzed using Perseus (Tyanova, Temu et al. 2016).

### Worm preparation for heat avoidance assay and number of replicates

Gravid adult worms were treated with hypochlorite solution according to standard protocol. Embryos were rinsed twice with water and once with M9 buffer, then resuspended in M9 and plated on NGM plates seeded with OP-50 *E. coli*. 200-300 embryos per individual plate were seeded. Worms were incubated at 20°C until start of egg laying (65-90 h, depending on the strain or condition). Worms were washed off the plates with distilled water, placed into 1.5 ml tubes and washed twice more to remove bacterial residues. Worms were placed on unseeded NGM plates and left to disperse and acclimate in the experimental room for 60 min. Prior to worm deposition, these experimental unseeded plates were kept open in a laminar flow hood for 3 h to ensure dry surface. Plate lid was removed 3 min before starting the assay. At least 3 replicates were performed for each strain or condition on 3 separate days running wild-type N2 strain alongside.

For the experiments using transgenic animals, fluorescent signal-carrying worms were picked 16-19 h before the experiment to allow for recovery.

### Heat stimulation and adaptation protocols

For the majority of experiments, we compared the heat-evoked response in naïve animals (that had never been stimulated, T0) and the same animals after 60 minutes of repeated stimulation (4-s stimuli and 20s interstimulus interval (ISI), T60). For some experiments, the adaptation period was reduced to 10 min (T10) or 30 min (T30). The INFERNO system (Lia and Glauser 2020) was used for behavioral recordings. The heat stimulation program during recordings was composed of a 40-s baseline period without any heat stimulation, a 4-s stimulation with 400 W heating (4 IR lamps turned on) and a 20-s post stimulation period. The repeated stimulation delivery (adaptation treatment) was achieved by placing the worm plates under the ThermINATOR system (Lia and Glauser 2020), providing infinitely looping temperature program composed of 4 s of IR lamps stimulation, followed by 20 s ISI with the lamps turned off. There was a lag of about 10 s for the plate transfer between the two systems.

In the INFERNO system, worm plates were recorded using a DMK 33U×250 camera and movies acquired with the IC capture software (The Imaging Source), at 8 frames per second, at a 1600×1800 pixel resolution, and the resulting .AVI file was encoded as Y800 8-bit monochrome. The Multi-Worm Tracker 1.3.0 (MWT) (Swierczek, Giles et al. 2011) with previously described configuration settings (Lia and Glauser 2020) was used for movie analyses. A previously described Python script was use to flag the frame of reversal occurrence (Lia and Glauser 2020). Each reported data point corresponds to the results of one assay plate scoring at least 50 animals.

### Transgene construction and transgenesis

Promoter-containing Entry plasmids (Multi-site Gateway slot 1) were constructed by PCR using N2 genomic DNA as template and primers flanked by attB4 and attB1r recombination sites; the PCR product being cloned into pDONR-P4-P1R vector (Invitrogen) by BP recombination.

Coding sequence-containing Entry plasmids (Multi-site Gateway slot 2) were constructed by PCR using N2 cDNA as template and primers flanked by attB1 and attB2 recombination sites; the PCR product being cloned into pDONR_221 vector (Invitrogen) by BP recombination.

Expression plasmids for transgenesis were created through a LR recombination reaction (Gateway^TM^ LR Clonase, Invitrogen) as per the manufacturer’s instructions.

Primer sequences for BP reactions and all plasmids used in this study are listed in File S4.

DNA constructs were transformed into competent DH5a *E. coli* (NEB C2987H), purified with the GenElute HP Plasmid miniprep kit (Sigma) and microinjected in the worm gonad according to a standard protocol (Evans 2006) together with co-injection markers for transgenic animal identification. The concentrations of expression plasmids and co-markers injected are indicated in File S4.

### Site-directed mutagenesis

To create truncated *tax-6(gf)* transgene PCR-based site-directed mutagenesis (Hemsley, Arnheim et al. 1989) was used. Entry plasmid containing *tax-6* coding DNA sequence was amplified with the KOD Hot Start DNA Polymerase (Novagen; Merck). Primers were designed to contain the desired deletion and phosphorylated in 5′ end (File S4). Linear PCR products were purified from agarose gel (1%) after electrophoresis with a Zymoclean-Gel DNA Recovery kit (Zymo Research), circularized using DNA Ligation Kit <Mighty Mix> (Takara).

### Statistical analyses

ANOVAs were conducted using Jamovi (The jamovi project (2022), jamovi (Version 2.3) [Computer Software]; retrieved from (https://www.jamovi.org). Post hoc tests were used to compare each of the mutants with wild type (N2) or *cmk-1(lf)* using Bonferroni–Holm correction.

## DATA AVAILABILITY

MS data have been deposited to the ProteomeXchange Consortium via the PRIDE partner repository with the dataset identifier PXD055776 (Perez-Riverol, Bai et al. 2022). The raw data and *p* values presented in the different figures are provided in File S3.

## ACKNOWLEDGEMENTS

We are grateful to Lisa Schild, Laurence Bulliard and Michael Stumpe for expert technical support. Some strains were provided by the CGC, which is funded by NIH Office of Research Infrastructure Programs (P40 OD010440). The study was supported by the Swiss National Science Foundation (BSSGI0_155764, PP00P3_150681, and 310030_197607 to DAG, 310030_212187 to JD) and by the Novartis Foundation for Medical-Biological Research.

## COMPETING INTERESTS

No competing interests declared

---

**Figure 1 -supplement 1.**
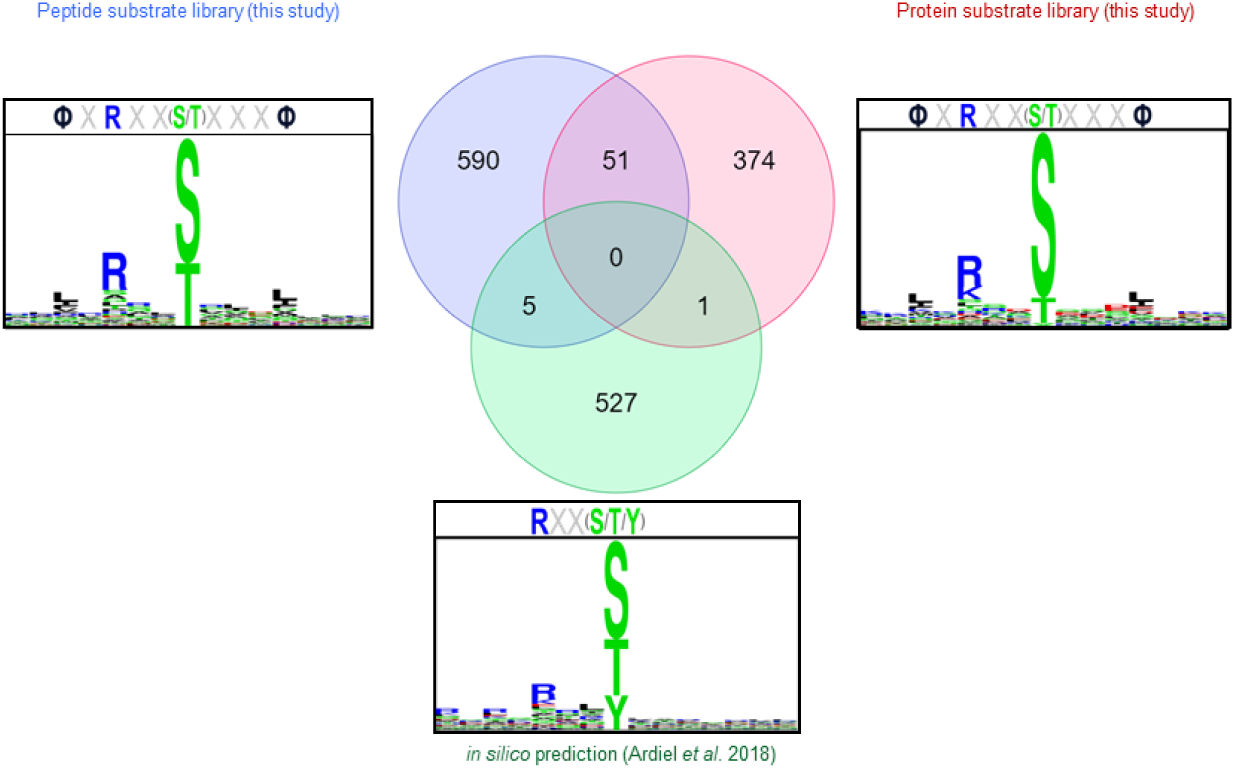
Comparison of empirically identified CMK-1 phospho-substrates with previously published predictions. Venn diagram showing the number of phosphosites from the indicated lists and the consensus as in Fig. 1.

**Figure 2-supplement 1.**
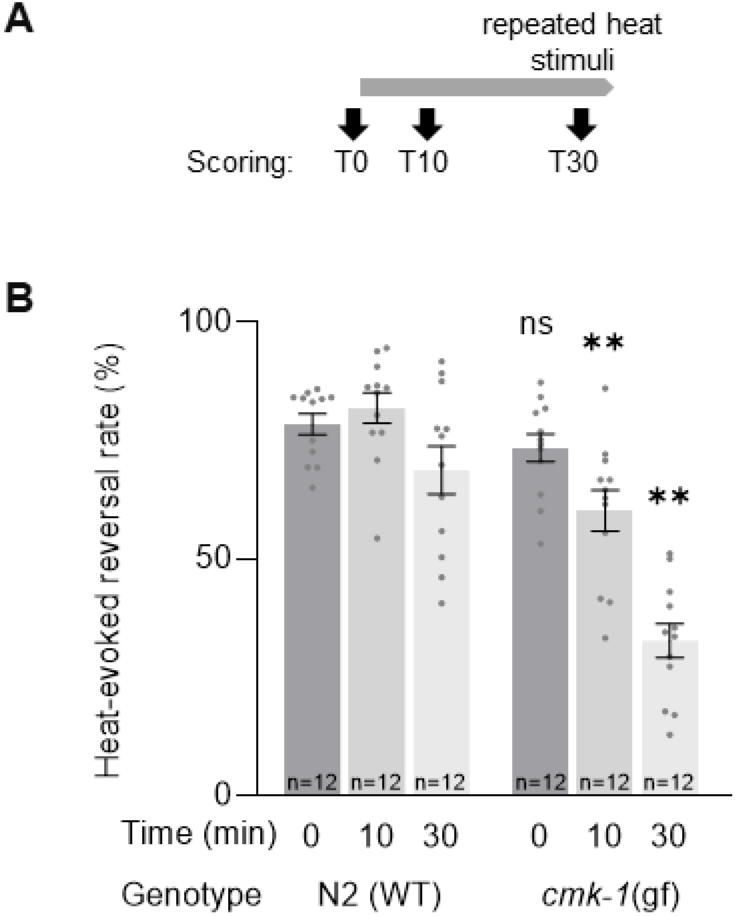
CMK-1 overactivating mutation T179D accelerates thermo-nociceptive adaptation. **A**. Schematic of the scoring procedure variation, including earlier scoring timepoints after 10 and 30 min of repeated noxious heat stimuli. **B**. Heat-evoked reversal scored in the indicated genotypes and reported as in Fig. 2. *cmk-1(gf)* is *cmk-1(syb1633)* harboring the CMK-1(T179D) overactivating mutation. *, *p<*.01 versus corresponding timepoint in N2(WT) control. ns, not significant.

**Figure 2-supplement 2.**
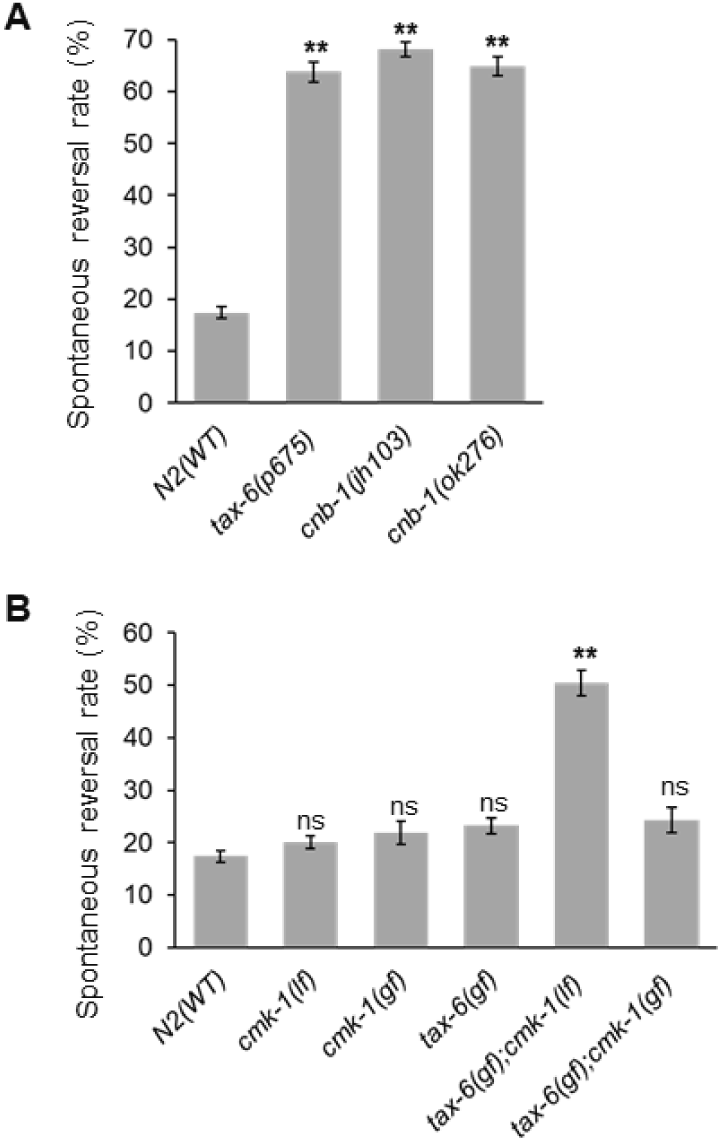
Spontaneous reversal rate in wild type and different single and double mutants. Spontaneous reversal rate scored in the indicated genotypes, in the absence of any noxious heat stimulation. Average (grey bars) and s.e.m (error bars) of *n≥*12 assays; each assay scoring at least 50 animals. *, *p<*.001 N2(WT) control. ns, not significant. **A**. Markedly elevated spontaneous reversal rate in *tax-6* and *cnb-1* loss-of-function mutants. **B**. Markedly elevated spontaneous reversal rate in *tax-6(gf);cmk-1(gf)* double mutant, but normal phenotype in other genotypes, as indicated.

**Figure 3- supplement 1.**
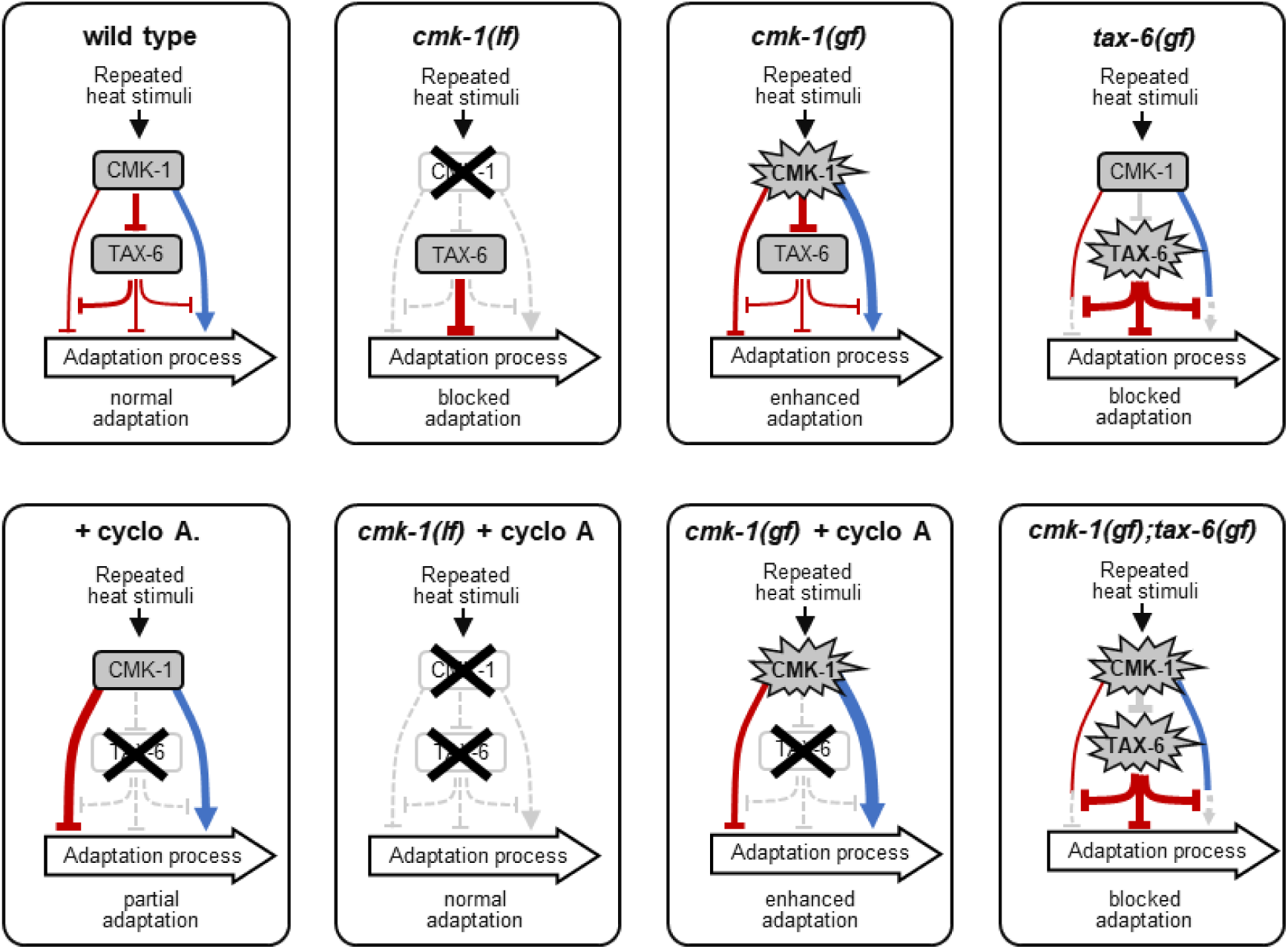
Model of the multiple antagonistic interactions observed between CMK-1 and TAX-6/CnA signaling. Illustration of how the model from Fig. 3D is proposed to be altered for each of the single or combined gain- and loss-of-function manipulations used in Fig. 3. We do not rule out that alternative models might also explain our data, but the model presented in Fig. 3D is the simplest that could explain the complex set of interactions. Blue regular arrows and red flathead arrows indicate positive and negative effects, respectively. Inactivated pathways are drawn with grey dashed lines. The thickness of each line reflects the relative size of an effect. The main assumption of the model is that CMK-1 and TAX-6 are modulators of the adaptation process and the balance between antagonistic effects will determine the level of adaptation. The default adaptation level in the absence of regulation (*cmk-1(lf)* + cyclo A condition) leads to a similar level of adaptation than in wild type. Furthermore, the model considers that the inhibitory effect of TAX-6 on the CMK-1 anti-adaptation branch and the inhibitory effect of TAX-6 on the CMK-1 pro-adaptation branch are not of the same magnitude. Likewise, the two antagonistic direct effects of CMK-1 on adaptation are not of the same magnitude: upon CMK-1 overactivation (*cmk-1(gf)*) the direct pro-adaptation effect of CMK-1 signaling is larger than the anti-adaptation effect. Finally, TAX-6 overactivation has a dominant effect (two panels at the right). cyclo A: Cyclosporin A treatment. *cmk-1(gf)* is *cmk-1(syb1633)* harboring the CMK-1(T179D) overactivating mutation. *cmk-1(lf)* is *cmk-1(ok287)*. *tax-6(gf)* is *tax-6(jh107)*.

## Notes

### Competing Interest Statement

The authors have declared no competing interest.

### Summary of Updates

The manuscript has been revised following evaluations and suggestions by peer reviewers.

