## Supplementary material for "Antagonist actions of CMK-1/CaMKI and TAX-6/Calcineurin along the *C. elegans* thermal avoidance circuit orchestrate adaptation of nociceptive response to repeated stimuli": File S4

**Strain list**

| **Strain Name** | **Genotype** | **Comments** |
|---|---|---|
| N2 | Wild type | Wild type (WT) |
| DAG800 | *cmk-1(pg58) IV* | Created previously (Schild et. al 2014) |
| DAG821 | *cmk-1(ok287) IV* | Obtained from CGC and then outcrossed twice with N2 |
| KJ306 | *tax-6 (jh107) IV* | Obtained from CGC. |
| PR675 | *tax-6 (p675) IV* | Obtained from CGC. |
| DAG712-713 | *cmk-1(ok287) IV; domEx712-713[gcy-5p:cmk-1:SL2::mCherry 20ng/ul; unc-122p::GFP 20ng/ul]* | *cmk-1* expression with *gcy-5* promoter driving *cmk-1* coding sequence (cds) in ASER neuron in *cmk-1* null background. |
| DAG716-717 | *cmk-1(ok287) IV; domEx716-717[gcy-6p:cmk-1:SL2::mCherry 20ng/ul; unc-122p::GFP 20ng/ul]* | *cmk-1* expression with *gcy-6* promoter driving *cmk-1* coding sequence (cds) in ASEL neuron in *cmk-1* null background. |
| DAG720-721 | *cmk-1(ok287) IV; domEx720-721[gcy-5p:cmk-1:SL2::mCherry 20ng/ul; gcy-6p::cmk-1::SL2mCherry 20ng/ul; unc-122p::GFP 20ng/ul]* | *cmk-1* expression with *gcy-5* and *gcy-6* promoters driving *cmk-1* coding sequence (cds) in ASER and ASEL neurons in *cmk-1* null background. |
| DAG819 | *cmk-1(ok287) IV, domEx 819 [mec-3p:cmk-1:SL2::GFP 20ng/ul, unc-122p::RFP 20ng/ul]* | *cmk-1* expression with *mec-3* promoter driving *cmk-1* coding sequence (cds) in FLP neurons in *cmk-1* null background. |
| DAG1002 | *cmk-1(syb1633) IV* | T179D substitution mutation made by genome editing (SunyBiotech, China). outcrossed 2x |
| DAG1405 | *cmk-1(ok287) IV; tax-6 (jh107) IV* | Obtained by crossing DAG821 and KJ307 |
| DAG1642-1644 | *domEx1642-1644[gcy-8p::QF, QUASp::TeTx::SL2mcherry, unc122p::GFP]* | Tetanus toxin expression with gcy-8 promoter in AFD neurons. |
| DAG1674-1676 | *cmk-1(ok287) IV [cmk-1p:cmk-1:SL2::GFP 20ng/ul, unc-122p::RFP 20ng/ul]* | *cmk-1* expression with *cmk-1* promoter driving *cmk-1* coding sequence (cds) in *cmk-1* null background. |
| DAG1708-1709 | *cmk-1(ok287) IV [tax-4p:cmk-1:SL2::GFP 20ng/ul, unc-122p::RFP 20ng/ul]* | *cmk-1* expression with *tax-4* promoter driving *cmk-1* coding sequence (cds) in sensory neurons in *cmk-1* null background. |
| DAG1710-1711 | *cmk-1(ok287) IV [glr-1p:cmk-1:SL2::GFP 20ng/ul, unc-122p::RFP 20ng/ul]* | *cmk-1* expression with *glr-1* promoter driving *cmk-1* coding sequence (cds) in interneurons in *cmk-1* null background. |
| DAG1765-1766 | *cmk-1(ok287) IV, domEx1765-1766 [str-2P::QF 20ng/ul, srsx-3P::QF 60ng/ul, QUAS::cmk-1::SL2mCherry, 20ng/ul, unc-122p::GFP 10ng/ul]* | *cmk-1* expression with *str-2* and *srsx-3* promoters driving *cmk-1* coding sequence (cds) in AWC neurons in *cmk-1* null background. |
| DAG1823 | *cmk-1(syb1633) IV; tax-6 (jh107) IV* | Obtained by crossing of DAG1002 and KJ307 |
| DAG1855-1857 | *domEx1855-1857[glr-1p::QF 20ng/ul, QUAS:tax-6(jh107):SL2::mCherry, 20ng/ul, unc-122p::GFP 10ng/ul]* | *tax-6(jh107)* truncated protein expression with *glr-1* promoter driving *tax-6* coding sequence (cds) in interneurons in WT background. |
| DAG1858-1859 | *domEx1858-1859[tax-4p::QF 20ng/ul, QUAS:tax-6(jh107):SL2::mCherry, 20ng/ul, unc-122p::GFP 10ng/ul]* | *tax-6(jh107)* truncated protein expression with *tax-4* promoter driving *tax-6* coding sequence (cds) in sensory neurons in WT background. |
| DAG1882-1884 | *domEx1888-1890[tdc-1::QF 20ng/ul, QUAS:tax-6(jh107):SL2::mCherry, 20ng/ul, unc-122p::GFP 20ng/ul]* | *tax-6(jh107)* truncated protein expression with *nmr-1* promoter driving *tax-6* coding sequence (cds) in RIM neurons in WT background. |
| DAG1885-1887 | *domEx1885-1887[cex-1::QF 20ng/ul, QUAS:tax-6(jh107):SL2::mCherry, 20ng/ul, unc-122p::GFP 20ng/ul]* | *tax-6(jh107)* truncated protein expression with *cex-1* promoter driving *tax-6* coding sequence |

| | | |
|---|---|---|
| | | (cds) in AVD and RIM neurons in WT background. |
| DAG1888-1890 | *domEx1888-1890[nmr-1::QF 20ng/ul, QUAS:tax-6(jh107):SL2::mCherry, 20ng/ul, unc-122p::GFP 20ng/ul]* | *tax-6(jh107)* truncated protein expression with *nmr-1* promoter driving *tax-6* coding sequence (cds) in AVD neurons in WT background. |
| DAG1868, DAG1968 DAG1969 | *cmk-1(ok287) IV; domEx1868,1968,1969[ttx-1p::QF 5ng/ul, QUAS:cmk-1:SL2::mCherry, 10ng/ul, unc-122p::GFP 10ng/ul]* | *cmk-1* expression with *ttx-1* promoter driving *cmk-1* coding sequence (cds) in AFD neuron in *cmk-1* null background. |
| DAG1932-1934 | *domEx1932-1934[mec-3p::QF 20ng/ul, QUAS:tax-6(jh107):SL2::mCherry, 20ng/ul, unc-122p::GFP 10ng/ul]* | *tax-6(jh107)* truncated protein expression with *mec-3* promoter driving *tax-6* coding sequence (cds) in FLP neurons in WT background. |
| DAG1935-1937 | *domEx1935-1937[lgc-39::QF 20ng/ul, QUAS:tax-6(jh107):SL2::mCherry, 20ng/ul, unc-122p::GFP 10ng/ul]* | *tax-6(jh107)* truncated protein expression with *lgc-39* promoter driving *tax-6* coding sequence (cds) in AVA neurons in WT background. |
| DAG1909-1910, 1913 | *cmk-1(ok287) IV; domEx1909-910,1913[gpa-4p::QF 20ng/ul, QUAS:cmk-1:SL2::mCherry, 20ng/ul, unc-122p::GFP 10ng/ul]* | *cmk-1* expression with *gpa-4* promoter driving *cmk-1* coding sequence (cds) in ASI neuron in *cmk-1* null background. |
| DAG1970-1972 | *cmk-1(ok287) IV; domEx1970-1972[ttx-1p::QF 5ng/ul, gcy-5p::QF 5ng/ul, QUAS:cmk-1:SL2::mCherry, 10ng/ul, unc-122p::GFP 10ng/ul]* | *cmk-1* expression with *ttx-1* and *gcy-5* promoters driving *cmk-1* coding sequence (cds) in AFD and ASER neurons in *cmk-1* null background. |
| GN112 | *pgIs2 [gcy-8p::TU#813 + gcy-8p::TU#814 + unc-122p::GFP + gcy-8p::mCherry + gcy-8p::GFP + ttx-3p::GFP]* | AFD ablation (gift from Miriam B. Goodman). |

| | | |
|---|---|---|
| **Promoter plasmids (multisitegateway slot 1)** | | |
| dg22 | Slot1 *Entry tax-4p* | Gift from Kaveh Ashrafi (UCSF, CA, USA) |
| dg25 | Slot1 *Entry glr-1p* | Gift from Kaveh Ashrafi (UCSF, CA, USA) |
| dg68 | Slot1 *Entry mec-3p(no ATG)* | Created previously (Schild et. al 2014) |
| dg229 | Slot1 *Entry QUASp* | Created previously (Schild et. al 2014) |
| dg507 | Slot1 *Entry gcy-8p* | Created previously (Ippolito et al. 2021) |
| dg508 | Slot1 *Entry ttx-1p* | Created previously (Ippolito et al. 2021) |
| dg763 | Slot1 Entry *gcy-5p* | Previously described (Lim et al. 2018) |
| dg764 | Slot1 Entry *gcy-6p* | Previously described (Lim et. al 2018) |
| dg833 | Slot1 *Entry gpa-4p* | Gift from Kaveh Ashrafi (UCSF, CA, USA) |
| dg949 | Slot1 *Entry srsx-3p* | Created previously (Jordan A, unpublished data) |
| dg950 | Slot1 *Entry str-2p* | Created previously (Jordan A, unpublished data) |
| dg1015 | Slot1 *Entry lgc-39p* | Created previously (Thapliyal et al. 2023) |
| dg1075 | Slot1 Entry *cex-1p* | Gift from Kaveh Ashrafi (UCSF, CA, USA) |
| dg1076 | Slot1 Entry *nmr-1p* | Gift from Kaveh Ashrafi (UCSF, CA, USA) |
| dg1077 | Slot1 Entry *tdc-1p* | Gift from Kaveh Ashrafi (UCSF, CA, USA) |
| mg237 | Slot1 *Entry mec-3p(w ATG)* | Created previously (Schild et. al 2014) |
| mg267 | Slot1 *Entry cmk-1p* | Created previously (Schild et. al 2014) |
| **Coding sequence plasmids (multisitegateway slot 2)** | | |
| dg88 | Slot2 Entry *tetx cds* | |
| dg240 | Slot2 Entry *QF* | Created previously (Schild et. al 2014) |
| dg335 | Slot2 Entry *cmk-1(K52A)cds* | Created previously (Schild et. al 2014) |
| dg592 | Slot2 Entry *cmk-1(T179D)cds* | Created previously (Schild et. al 2014) |
| dg725 | Slot2 Entry *tax-6cds* | Generated by BP reaction using mg205 |
| | Primers:<br>attB1-Tax-6_F :<br>ggggacaagtttgtacaaaaaagcaggctTAATGGCCTCGACATCGGCAGGAC<br>attB2-Tax-6_R:<br>ggggaccactttgtacaagaaagctgggtCGCTATTTGATGGACCATTTTGTGG | |
| dg1031 | Slot2 Entry *tax-6(jh107)cds* | Generated by Site-directed mutagenesis from dg725 |
| | Primers:<br>tax-6_j107SDM_STOP_Fw: TTTAACTAAGACCCAGCTTTCTTGTACAAAG<br>tax-6_kj306SDM_Rw: AAAACCATTTCCAATTGCTCGAATCTTGTG | |
| mg269 | Slot2 Entry *cmk-1cds(no ATG)* | Created previously (Schild et. al 2014) |
| mg271 | Slot2 Entry *cmk-1cds(w ATG)* | Created previously (Schild et. al 2014) |
| **Expression plasmids used for transgenesis** | | |
| dg6 | *mec-3*p:*cmk-1*:SL2::GFP | Created through a LR recombination reaction between mg237, mg269, mg276, dg560 |
| dg7 | *cmk-1*p:*cmk-1*:SL2::GFP | Created through a LR recombination reaction between mg267, mg269, mg276, dg560 |
| dg28 | *tax-4*p:*cmk-1*:SL2::GFP | Created through a LR recombination reaction between dg22, mg269, mg276, dg560 |
| dg31 | *glr-1*p:*cmk-1*:SL2::GFP | Created through a LR recombination reaction between dg25, mg269, mg276, dg560 |
| dg243 | *mec-3p::QF::unc-54UTR* | Created through a LR recombination reaction between dg68, dg240, mg211, dg560 |

| | | |
|---|---|---|
| dg249 | *QUAS::cmk-1::SL2::mCherry* | Created through a LR recombination reaction between dg229, mg271, mg277, dg560 |
| dg255 | *QUAS::TeTx::SL2::mCherry* | Created through a LR recombination reaction between dg229, dg88, mg277, dg560 |
| dg616 | *gcy-5*p:*cmk-1*:SL2::mCherry | Created through a LR recombination reaction between dg763, mg271, mg277, dg560 |
| dg617 | *gcy-6*p:*cmk-1*:SL2::mCherry | Created through a LR recombination reaction between dg764, mg271, mg277, dg560 |
| dg783 | *srsx-3p*::QF::unc-54UTR | Created through a LR recombination reaction between dg949, dg240, mg211, dg560 |
| dg785 | *str-2p*::QF::unc-54UTR | Created through a LR recombination reaction between dg950, dg240, mg211, dg560 |
| dg845 | *gpa-4*p::QF::unc-54UTR | Created through a LR recombination reaction between dg833, dg240, mg211, dg560 |
| dg883 | *ttx-1*p::QF::unc-54UTR | Created through a LR recombination reaction between dg508, dg240, mg211, dg560 |
| dg931 | *gcy-8p::QF::*unc-54UTR | Created through a LR recombination reaction between dg507, dg240, mg211, dg560 |
| dg957 | *glr-1p::QF::SL2::mCherry* | Created through a LR recombination reaction between dg25, dg240, mg277, dg560 |
| dg1017 | *lgc-39p::QF::unc-54UTR* | Created through a LR recombination reaction between dg1015, dg240, mg211, dg560, |
| dg1026 | *tax-4p::QF::unc-54UTR* | Created through a LR recombination reaction between dg22, dg240, mg211, dg560 |
| dg1027 | *gcy-5*p::QF::unc-54UTR | Created through a LR recombination reaction between dg763, dg240, mg211, dg560 |
| dg1028 | *gcy-6*p::QF::unc-54UTR | Created through a LR recombination reaction between dg764, dg240, mg211, dg560 |
| dg1059 | *QUAS::tax-6(jh107)::SL2::mCherry* | Created through a LR recombination reaction between dg229, dg1031, mg277, dg560 |
| dg1078 | *cex-1p::QF::unc-54UTR* | Created through a LR recombination reaction between dg1075, dg240, mg211, dg560 |
| dg1079 | *nmr-1p::QF::unc-54UTR* | Created through a LR recombination reaction between dg1076, dg240, mg211, dg560 |
| dg1080 | *tdc-1p::QF::unc-54UTR* | Created through a LR recombination reaction between dg1077, dg240, mg211, dg560 |
| **3' UTR and tagging plasmids (multi-site gateway slot3)** | | |
| mg277 | *slot3 Entry SL2::mCherry* | Previously described (Schild et. al 2014) |
| mg211 | *slot3 Entry unc-54 3'UTR* | gift from Marc Hammarlund (Yale University, CT, USA) |
| mg276 | *slot3 Entry SL::GFP* | Created previously (Schild et. al 2014) |
| **Co-injection markers** | | |
| dg9 | *unc-122*p::RFP | gift from Piali Sengupta (Brandeis university, MA, USA); Addgene plasmid # 8938 |
| dg396 | *unc-122p::*GFP | gift from Piali Sengupta (Brandeis university, MA, USA); Addgene plasmid # 8937 |
| **DONR and DEST plasmids for vector construction** | | |
| mg169 | *pDONR P4 P1R* | gift from Marc Hammarlund (Yale University, CT, USA) |
| mg205 | *pDONR 221* | gift from Miriam B. Goodman (Stanford University, CA, USA) |
| dg560 | *pDEST R4-R3* | gift from Marc Hammarlund (Yale University, CT, USA) |

| Plasmids used for recombinant protein expression | | |
|---|---|---|
| dg773 | pET-24d/GST-TEV- cmk-1, 1-295, T179D | Was created by insertion of cmk-1 cds (from dg592) using NdeI and BamHI restriction sites in pDK2409 |
| dg776 | pET-24d/GST-TEV- cmk-1, 1-295, K52A | Was created by insertion of cmk-1 cds (from dg335) using NdeI and BamHI restriction sites in pDK2409 |
| dg728 | pET-24d/(His)6- tax-6 | Was created by insertion of tax-6 cds (from dg725) using NdeI and BamHI restriction sites in pDK2832 |
